# Museomics of *Carabus* giant ground beetles shows an Oligocene origin and *in situ* alpine diversification

**DOI:** 10.1101/2024.03.21.586057

**Authors:** Marie T. Pauli, Jérémy Gauthier, Marjorie Labédan, Mickael Blanc, Julia Bilat, Emmanuel F.A. Toussaint

## Abstract

The development of museomics represents a major paradigm shift in the use of natural history collection specimens for systematics and evolutionary biology. New approaches in this field allow the sequencing of hundreds to thousands of loci from across the genome using historical DNA. HyRAD-X, a recently introduced capture method using bench-top designed probes, has proved very efficient for recovering genomic-scale datasets using natural history collection specimens. Using this technique at both the intra- and interspecific levels, we infer the most robust phylogeny to date for Arcifera, an ecologically and morphologically diverse clade of *Carabus* giant ground beetles. We successfully generated a genomic dataset of up to 1965 HyRAD-X loci for all described species, permitting inference of a robust dated phylogenomic tree for this clade. Our species delimitation and population genomic analyses suggest that the current classification for Arcifera is in line with its evolutionary history. Our results suggest an origin of Arcifera in the late Oligocene followed by speciation events during the warm mid-Miocene unlinked to Pleistocene glaciations. The dynamic paleogeographic history of the Palearctic region likely contributed to the diversification of this lineage with a relatively ancient colonization of the proto-Alps followed by *in situ* speciation where most species of Arcifera are currently found sometimes syntopically likely as a result of post-glaciations secondary contacts.

## Introduction

Recent developments in museomics are opening new prospects allowing samples from natural history collections (NHC) to enter the era of genomics (reviewed in Raxworthy & Smith 2021; Card *et al*. 2021). Specimens held in the collections are crucial for the study of systematics and taxonomy, but also for the study of ecology and evolution (Duchenne *et al*. 2020). Sampling from NHC specimens is a major asset to study groups that are currently rare in the wild, for which authorizations to collect new specimens are difficult to obtain, or for which a comprehensive taxonomic and geographic sampling would require extensive fieldwork campaigns. Such a strategy is therefore very powerful when working on taxonomic groups presenting a wide geographical range. In extreme cases, and when species are believed to be extinct, NHC represent the only potential source of genetic data (Toussaint *et al*. 2021; de-Dios *et al*. 2023).

Innovative approaches are now making it possible to obtain genetic information from NHC specimens for which it has long been impossible to recover DNA. The DNA in these specimens, referred to as historical DNA (hDNA), is in low quantity, fragmented, has undergone chemical modifications over time and contains contaminants linked to the history of the collection (Raxworthy & Smith 2021). Improvements in extraction methods, sequencing technologies but above all the development of new capture methods have enabled an increasing amount of genetic information to be recovered. They allow difficulties associated with highly degraded and fragmented hDNA from NHC samples to be overcome, which prevent conventional amplification using standard molecular primers (Landry *et al*. 2023). Among these methods, Ultra Conserved Elements (Blaimer *et al*. 2016; Faircloth 2017) or anchored hybrid enrichment of conserved regions (AHE, Lemmon *et al*. 2012; Mayer *et al*. 2021), are based on the capture of informative loci previously designed from existing genomic data and generally target fairly conserved regions in order to produce large phylogenies. Applying these approaches to NHC specimens allows integration of samples that are complicated to obtain in the field. In order to work on non-model species for which no prior genomic data is available, the HyRAD (Suchan *et al*. 2016) and HyRAD-X (Schmid *et al*. 2017) approaches enable probes to be designed directly from a few phylogenetically close fresh samples. These approaches based on bench-top production are considerably less expensive than commercially synthesized probes. The probes are designed using a ddRADseq protocol (Peterson *et al*. 2012) to target thousands of loci randomly distributed along the genome. This approach is suitable for integrating NHC samples into population-scale studies (Gauthier *et al*. 2020) or for phylogenetic studies of taxa that have recently diverged, such as within a genus (Gauthier *et al*. 2023). The HyRAD-X approach designs probes on fresh RNA extractions. By targeting only expressed gene loci, the HyRAD-X approach makes it possible to investigate phylogenetic questions at older evolutionary scales than the HyRAD approach (Toussaint *et al*. 2021). Using these probe sets, hDNA is then captured by hybridization and sequenced using NGS technologies. This allows only the targeted loci to be recovered while eliminating all unwanted fragments such as contaminants. After sequencing, the loci are reconstructed and aligned using appropriate bioinformatic pipelines in order to make phylogenetic inferences (Toussaint *et al*. 2021). Unlike random Whole Genome Sequencing (WGS) of all the extracted DNA, these targeted approaches enable better recovery of loci and integration of a larger number of NHC samples into the phylogenetic inferences (Toussaint *et al*. 2021). Although the efficiency of HyRAD-X has been tested at higher taxonomic levels, an empirical investigation of its performance at the interface between population and species levels is needed.

The genus *Carabus*, Linnaeus 1758 (Coleoptera: Carabidae), is a monophyletic highly diversified lineage comprising *ca.* 970 species classified into 91 subgenera (Deuve 2019, 2021). This genus, together with its sister genus *Calosoma* (cosmopolitan, 130 species) form the tribe Carabini (Osawa *et al*. 2004; Toussaint Fls & Gillett 2018; Toussaint *et al*. 2021; Sota *et al*. 2022). Within *Carabus*, the clade named Arcifera Imura, 1996 is sister to the very diversified clade Eucarabi Deuve, 2013 (Deuve *et al*. 2012; Deuve 2021). This clade is mainly Palearctic, ranging in the west from southwest England to Ukraine and Turkey in the east. The range of this group notably encompasses the Carpathian Mountains as well as the Swiss, Italian, Austrian and Dinaric Alps. It currently includes four subgenera: *Carabus (Hygrocarabus*) Thomson, 1875, *Carabus (Platycarabus)* Morawitz, 1886, *Carabus (Chaetocarabus*) Thomson, 1875 and *Carabus (Heterocarabus)* Morawitz, 1886 (Deuve 2019, 2021).

Within Arcifera, the subgenus *C. (Hygrocarabus*) contains two species, *Carabus nodulosus* Creutzer, 1799 and *Carabus variolosus* Fabricius, 1787, found from France to Ukraine, the status of which has been extensively debated over the past decades due to reduced morphological differences and inconsistent genetic admixture patterns (Müller-Kroehling *et al*. 2006, 2014; Matern *et al*. 2009, 2010; Mossakowski *et al*. 2020). These hygrophilous nocturnal species live in river banks and hunt close to or in the water of cold forest streams. The two species are in relative allopatry with *C. nodulosus* being found from eastern France to Austria and western Balkans, and *C. variolosus* from Slovakia to Ukraine and Bulgaria (Kulijer 2019; Deuve 2021; Bekchiev *et al*. 2022; Hristovski *et al*. 2023). Despite the protection status of their habitat (Annexes II and IV of the European Union’s Habitats Directive), these two species appear to be declining due to anthropogenic activities and their consequences (Tyszecka *et al*. 2023).

The subgenus *C.* (*Chaetocarabus*) only contains two allopatric species following Deuve (Deuve 2019, 2021), the widespread *Carabus intricatus* Linné, 1761 found from western France to Ukraine and Greece, and the Greek endemic *Carabus arcadicus* Gistl, 1850. These two species are found in sympatry in Greece where hybrids are known (at the Katara Pass in the Epirus region for instance) (Arndt et al. 2011). Additionally, the status of several subspecies in both taxa has been debated, and some authors recognize *Carabus arcadicus merlini* Schaum, 1861 (Greece), *Carabus intricatus lefebvrei* Dejean, 1826 (southern Italy including Sicily) and *Carabus intricatus krueperi* Reitter, 1896 (Greece) as separate species within which additional subspecific taxa have been described (e.g., Cavazzuti & Ghiretti 2020). Perhaps the most debated taxon of the three is *Carabus intricatus lefebvrei* found south of Umbria to northern Sicily, which is largely allopatric from the rest of the Italian populations found only in the extreme north of Italy from Piemonte to Friuli (Cavazzuti & Ghiretti 2020).

The subgenus *C. (Heterocarabus)* contains a unique species, *Carabus marietti* Cristofori & Jan, 1837, that is found in southern Bulgaria near the Black Sea and in Anatolia (Turkey). However the ecology and relationships between its numerous described subspecies remain poorly known (Gueorguiev & Gueorguiev 1995; Hieke & Wrase 2008).

Finally, the subgenus *C. (Platycarabus)* is composed of five currently accepted species: *Carabus creutzeri* Fabricius, 1801, *Carabus cychroides* Baudi, 1860, *Carabus depressus* Bonelli, 1811, *Carabus fabricii* Panzer, 1812 and *Carabus irregularis* Fabricius, 1792. These beetles are characterized by a flattened morphology, long legs, and elytra generally covered with small foveae (except in *Carabus depressus lucens* Schaum, 1857). They are most widely distributed in Central and Eastern Europe, generally at high altitudes, in mountain forests and alpine pastures. The subgenus contains helicophagous species that exhibit different hunting techniques related to the morphology of their mandibles and prothorax (Casale *et al*. 1998). For instance, *Carabus cychroides* has undergone a process known as “cychrization” (Thiele 1977; Casale *et al*. 1998; Symondson 2004), a process by which the pronotum is narrowed to allow predation inside snail shells (= stenocephalic morphology) as observed for instance in members of the tribe Cychrini. This species has a very restricted range in the Piedmont region of Italy, is endangered and the focus of reinforced conservation programs (Anselmo & Rizzioli 2022a; b). In contrast, the species *C. irregularis* presents a “licinization” or “procerization” process (Thiele 1977; Casale *et al*. 1998; Symondson 2004) with a macrocephalic morphology displayed by a large head adapted to cracking snail shells. The relationships among species of the subgenus *C.* (*Platycarabus*) are still debated, and the various taxonomic divisions, both species and subspecies, have yet to be clarified (Casale *et al*. 1998; Deuve 2021). Natural hybrids have been suggested between *C. fabricii* and *C. depressus*, *C. creutzeri* and *C. irregularis*, *C. creutzeri* and *C. depressus*, and *C. depressus* and *C. cychroides* (Casale *et al*. 1998; Camard & Leplat 2004; Casale & Rapuzzi 2015), indicating the need for an in-depth study of possible hybridization in this group.

One of the earliest attempts to elucidate the phylogeny of Arcifera was conducted by Ishikawa (Ishikawa 1984), using 21 morphological characters. This study supported the monophyly of Arcifera and placed *C. (Hygrocarabus*) as sister to the rest of the group, in which *C.* (*Chaetocarabus*) was sister to *C. (Heterocarabus)* and *C. (Platycarabus)*. The first placement of Arcifera members in a molecular phylogeny was based on a single mitochondrial fragment (i.e. ND5), and recovered *C.* (*Chaetocarabus*) and *C. (Platycarabus)* as sister lineages, close to *Carabus* (*Limnocarabus*) Géhin, 1876 and *Carabus* (*Euleptocarabus*) Nakane, 1956 (Imura *et al*. 1998). A subsequent study with the same gene fragment but increased taxon sampling recovered a paraphyletic Arcifera due to the placement of *C. (Hygrocarabus*) as sister to *Carabus* (*Limnocarabus*) and *C.* (*Euleptocarabus*) (Imura *et al*. 2000). In the same study, *C. (Heterocarabus)* was sister to *C.* (*Chaetocarabus*) and *C. (Platycarabus)*. Using the same gene fragment, another study inferred *C. (Platycarabus)* as sister to *C.* (*Chaetocarabus*) and *C. (Heterocarabus)*, within a largely unresolved *Carabus* clade (Su *et al*. 2003). A subsequent study using two nuclear gene fragments recovered Arcifera, represented by *C.* (*Chaetocarabus*) and *C. (Platycarabus)* as sister to the rest of the genus (=Eucarabi) (Sota & Ishikawa 2004). More recently, Deuve *et al*. (2012) used ten loci to recover Arcifera as sister to the Eucarabi and within Arcifera, they recovered *C. (Hygrocarabus*) as sister to *C.* (*Chaetocarabus*) and *C. (Platycarabus)*.

Phylogenetic relationships among *C. (Platycarabus)* species were also investigated using Sanger sequencing data (Casale *et al*. 1998), suggesting that *C. irregularis* is sister to the rest of the subgenus with *C. fabricii* and *C. depressus* being the most derived lineages in the tree. In parallel to a moderate refinement in the phylogenetic inferences of Arcifera, the estimation of divergence times in the clade has made some progress. Estimates for the origin of Arcifera based on few loci range from the mid-Miocene (ca. 14 Ma, Deuve *et al*. 2012) to the early Oligocene (ca. 30 Ma, Schmidt *et al*. 2023). No major improvement in our understanding of Arcifera systematics and evolution has been achieved in the past decade and there is a need to infer a robust evolutionary tree for this section of *Carabus* to better understand the morphological, ecological, and geographical evolution of constituent lineages.

In this study, we take advantage of the HyRAD-X approach to integrate a large number of samples throughout the geographical range of Arcifera. We rely on phylogenomic inferences, species delimitations and population genomics approaches to clarify the taxonomy and elucidate the evolutionary history of this complex group of species. In particular, we use this new genomic framework to test which abiotic factors may have fostered the diversification of Arcifera through space and time in the Cenozoic.

## Material and methods

### Taxon sampling and DNA extraction

The initial sampling was designed in order to sample major lineages within the Arcifera group comprising four subgenera *C.* (*Chaetocarabus*), *C. (Heterocarabus)*, *C. (Hygrocarabus*) and *C. (Platycarabus)* (Deuve 2019). A total of 96 samples were initially collected, mainly from NHC samples (87 samples, i.e. 90% of the dataset) but also from a few fresh samples (9 samples, i.e. 10% of the dataset) when these were available (Supplementary Table 1). Multiple specimens of the same taxa and geographic populations were initially selected to anticipate the risk of failure linked to hDNA degradation that can result in specimens being excluded. NHC specimens used in this study are kept at the Natural History Museum of Geneva (MHNG, 76 specimens) and Zoologische Staatssammlung München (ZSM-SNSB, 10 specimens). Eight specimens collected in 96% ethanol were also used and have been deposited in the MHNG collections. DNA was extracted destructively from a single leg using a QIAamp DNA Micro kit (Qiagen, Hilden, Germany) and eluted in 20µL of ultrapure water. Quantity and quality of the purified DNA were assessed with a Fragment Analyzer. Based on DNA quality and concentrations, 38 specimens were not included in the final samples selected for capture, enrichment and sequencing (*ca.* 40% of DNA extractions not processed). Overall, a total of 56 Arcifera specimens were sequenced de novo for this study, representing all Arcifera subgenera and species, several subspecies for the most widespread species as well as good geographical representation of each species range (taking into account 12 specimens that were eventually not included in the decisive datasets, see Results).

Early sampling erosion and discarded samples are commonly not discussed in the framework of museomics studies, but we believe that this is critical to understand the limitations and cost of such approaches in modern phylogenomic studies. The initial sampling in this study was specifically designed to accommodate a *ca.* 40–50% specimen loss during DNA quality/quantity assessment (e.g., Toussaint *et al*. 2021), and therefore the resulting sampling is well-suited to tackle the focal taxonomic and evolutionary questions in Arcifera. The final taxon sampling was complemented by eight samples of *Carabus* (including one of *C. irregularis* and one of *C. variolosus*) and one of *Calosoma sycophanta* (Linné, 1758) retrieved from (Toussaint *et al*. 2021) (see Supplementary Table 1 for more details).

### HyRAD-X protocol

The HyRAD protocol was applied as in (Toussaint *et al*. 2021) allowing the same probe set to be generated and therefore backward compatibility with the data of this previous study. For fresh specimens a shearing step with NEBNext dsDNA Fragmentase (New England Biolabs) was performed before library preparation. Shotgun libraries were prepared based on the protocol developed in Tin *et al*. (2014). Purified DNA was phosphorylated with T4 Polynucleotide Kinase. After heat-denaturation into single-stranded DNA, G-tailing was performed with Terminal Transferase and second strand DNA was synthesized with Klenow Fragment (3 ->5 exo-) using a poly-C oligonucleotide. Blunt-end reaction was performed with T4 DNA Polymerase and barcoded adapters were ligated to the phosphorylated end with T4 DNA ligase. After adapter fill-in with Bst DNA Polymerase (Large Fragment), PCRs were run using Phusion U Hot Start DNA Polymerase (Thermo Scientific) and indexed PCR primers. Libraries were pooled in equimolar quantities based upon their respective concentrations. Hybridization capture for enrichment of shotgun libraries was based on the MYbaits protocol (Arbor Biosciences) modified as in (Toussaint *et al*. 2021) to include a two-step capture at different temperatures (Li *et al*. 2013). Final library sequencing was performed on Illumina NovaSeq 6000 SP using a paired-end protocol (Lausanne Genomic Technologies Facility, Switzerland).

### Illumina sequencing data cleanup and processing

Raw reads were demultiplexed according to indexes and barcodes using CutAdapt2 (Martin 2011). Reads were cleaned using CutAdapt2 (Martin 2011) and quality was assessed all along the process using fastqc (https://www.bioinformatics.babraham.ac.uk/projects/fastqc/). Cleaned reads were individually mapped onto the loci catalog using BWA-MEM (Li 2013) (Supplementary Figure 1). The GATK (GenomeAnalysisTK) IndelRealigner tool (McKenna *et al*. 2010) realigned the indels and deamination were corrected using *mapDamage2.0* (Jónsson *et al*. 2013). For each sample and each locus, a consensus sequence was generated from the mapping file using samtools mpileup, bcftools and *vcfutils.pl* (Li *et al*. 2009). Consensuses were generated keeping the majority allele at each position. Twelve samples with too much missing data (more than 80% of N), were identified using seqtk and removed (Supplementary Table 1). Two thresholds of minimum coverage (min_cov) were applied to keep positions: min_cov=3 and min_cov=6. To test different levels of missing data, decisive datasets were generated applying three thresholds for the minimum number of samples per locus (min_sample): min_sample=10, min_sample=17 and min_sample=32. As a result, six datasets were generated: Dataset A (min_cov=6, min_sample=10, 50 taxa, 1,481 loci), Dataset B (min_cov=6, min_sample=10, 52 taxa, 1,965 loci), Dataset C (min_cov=6, min_sample=17, 50 taxa, 1,014 loci), Dataset D (min_cov=3, min_sample=17, 52 taxa, 1,291 loci), Dataset E (min_cov=6, min_sample=32, 50 taxa, 366 loci) and Dataset F (min_cov=3, min_sample=32, 52 taxa, 478 loci) (Table 1). The consensus sequences were combined and aligned with MAFFT using the --auto option. Eight samples from (Toussaint *et al*. 2021) were integrated at the alignment step. The final datasets only differ at the taxon sampling level with respect to *C. arcadicus merlini* CBX0176 and *C. cychroides* CBX0082 that were included only in Datasets B, D and F (these two taxa were systematically discarded because of low genomic coverage when generating loci with a min_cov=6, i.e., in Datasets A, C and E).

**Table 1.**
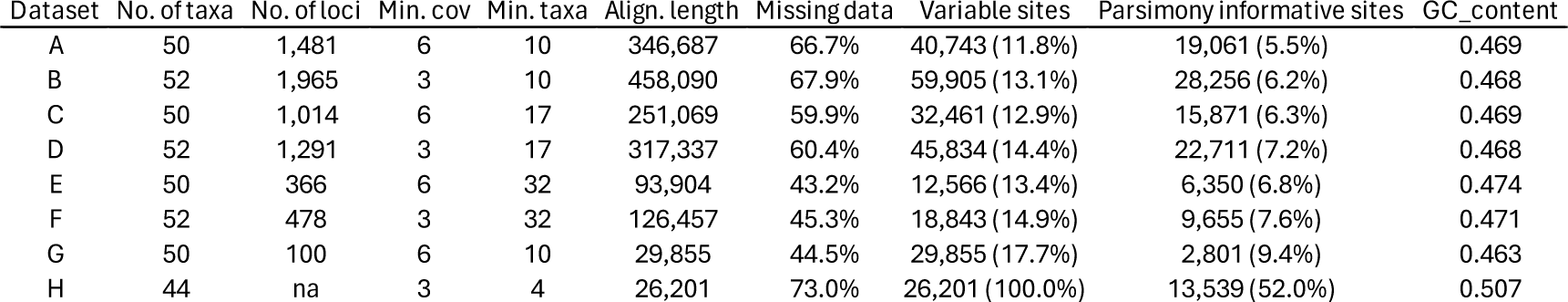
Alignment statistics for each dataset, including the number of taxa, the number of loci, the minimum coverage, the minimum number of taxa, the alignment length, the percentage of missing data, the numbers and percentages of variable sites and of parsimony informative sites, and the GC content.

| Dataset | No. of taxa | No. of loci | Min. cov | Min. taxa | Align. length | Missing data | Variable sites | Parsimony informative sites | GC_content |
| --- | --- | --- | --- | --- | --- | --- | --- | --- | --- |
| A | 50 | 1,481 | 6 | 10 | 346,687 | 66.7% | 40,743 (11.8%) | 19,061 (5.5%) | 0.469 |
| B | 52 | 1,965 | 3 | 10 | 458,090 | 67.9% | 59,905 (13.1%) | 28,256 (6.2%) | 0.468 |
| C | 50 | 1,014 | 6 | 17 | 251,069 | 59.9% | 32,461 (12.9%) | 15,871 (6.3%) | 0.469 |
| D | 52 | 1,291 | 3 | 17 | 317,337 | 60.4% | 45,834 (14.4%) | 22,711 (7.2%) | 0.468 |
| E | 50 | 366 | 6 | 32 | 93,904 | 43.2% | 12,566 (13.4%) | 6,350 (6.8%) | 0.474 |
| F | 52 | 478 | 3 | 32 | 126,457 | 45.3% | 18,843 (14.9%) | 9,655 (7.6%) | 0.471 |
| G | 50 | 100 | 6 | 10 | 29,855 | 44.5% | 29,855 (17.7%) | 2,801 (9.4%) | 0.463 |
| H | 44 | na | 3 | 4 | 26,201 | 73.0% | 26,201 (100.0%) | 13,539 (52.0%) | 0.507 |

MitoFinder (Allio *et al*. 2020) was used to identify mitochondrial genes among all sequenced loci. The kept genes were shared by at least half of the samples. The genes were aligned, and the sequences were cleaned in Geneious. Individual locus haplotype networks were built in SplitsTree v.4.19.1 (Huson & Bryant 2006). The networks were reconstructed using calculated uncorrected p-distances and the NeighborNet algorithm. All non-Arcifera outgroups were removed before analyses. In parallel, SNP calling was performed on the mapping files from the Arcifera species using GATK (McKenna *et al*. 2010) in order to perform complementary population genomic analyses and compare the results with those obtained from the locus-oriented approach, avoiding any bias linked to locus reconstruction (Dataset H).

### Phylogenetic inferences

For each dataset, phylogenetic inferences were performed using IQ-TREE v2.0.5 (Minh *et al*. 2020) using the edge-linked partition model (Chernomor *et al*. 2016). First, the best partitioning schemes were estimated using PartitionFinder v2.1.1 (Lanfear *et al*. 2017) with the rcluster algorithm under the Akaike Information Criterion corrected (AICc), with a rcluster-max of 2,000 and a rcluster-percent of 20. The resulting partitioning schemes were then used in IQ-TREE to select corresponding models of nucleotide substitution using ModelFinder (Kalyaanamoorthy *et al*. 2017) and the AICc across all available models in IQ-TREE. To avoid local optima, we performed 100 independent tree searches for each dataset in IQ-TREE. To estimate branch support, we calculated 1,000 ultrafast bootstraps along with 1,000 SH-aLRT tests in IQ-TREE (Guindon *et al*. 2010; Hoang *et al*. 2018). We used the hill-climbing nearest-neighbour interchange topology search strategy to avoid severe model violations leading to biased ultrafast bootstrap estimations (Hoang et al., 2018). The best tree for each analysis was selected based on the comparison of maximum likelihood scores. Coalescent species trees were inferred using ASTRAL-hybrid (Zhang & Mirarab 2022). We first performed individual locus trees using IQ-TREE v2.0.5 (Minh *et al*. 2020) and branch supports were assessed using 1,000 ultrafast bootstraps. Best substitution model for each locus was estimated using ModelFinder (Kalyaanamoorthy *et al*. 2017). Species tree reconstruction was performed combining gene trees using the weighted-ASTRAL optimization algorithm (Zhang & Mirarab 2022) taking into account phylogenetic uncertainty by relying on branch length and branch support across locus trees. As a complement to the locus reconstruction approach, we performed phylogenetic inferences based on the SNP set used for the population genomic approaches. Bi-allelic SNPs shared by at least four samples were extracted and all invariant sites removed. Species trees were inferred with RAxML-NG (Kozlov *et al*. 2019) using GTR+G+ASC_LEWIS model for ascertainment bias correction and branch supports were assessed using 1,000 bootstraps.

### Divergence time estimation

Divergence time estimation was performed in BEAST 1.10.4 (Suchard *et al*. 2018) based on a subset of loci selected using a gene-shopping approach to make these analyses tractable on a bioinformatic cluster. SortaDate (Smith *et al*. 2018) was used with default settings to select 100 loci sorted using the following criteria: clock-likeness, tree length, and least topological conflict with the IQ-TREE species tree on dataset E. The selected loci were then concatenated into a Dataset G for relaxed-clock Bayesian divergence time estimation. The best partitioning scheme and substitution models were determined with PartitionFinder2 (Lanfear *et al*. 2017) using the greedy algorithm with the parameter minsubset-size = 2000 and the Bayesian information criterion algorithm to choose between competing models. Clock partitioning was implemented by 1) a single clock for all partitions and 2) a clock for each partition (eight in total; see Results). A Bayesian lognormal relaxed clock model was assigned to the different clock partitions. Different tree models were tested using a Yule pure birth model (Yule 1925; Gernhard 2008), a birth-death model (Drummond *et al*. 2006; Gernhard 2008) as well as a Constant population size coalescent model (Kingman 1982). Since the fossil record of *Carabus* is scarce, we relied on secondary calibrations from a study focusing on Adephaga evolution based on 23 beetle fossil calibrations (Baca *et al*. 2021). According to this study, the separation between the genera *Calosoma* and *Carabus* occurred about 41.4 [37.1–46.1] million years ago (Ma). This age was used as a secondary calibration for the corresponding node in our topology (split *Calosoma*/*Carabus*, in this case the root). A second calibration was used to constrain the crown of *Carabus*. Following (Baca *et al*. 2021), this node was constrained to match the recovered age in their study at about 25.4 [22.8–28.2] Ma. The analyses were conducted for 50 million generations, sampling parameters and trees every 5000 generations. The maximum clade credibility tree for each analysis was generated in TreeAnnotator 1.10.4.

### Species delimitation and hybridization

We used a combination of species delimitation methods and population genomic approaches to test species and subspecies limits. For these analyses we excluded the six non-Arcifera outgroup specimens resulting in a dataset composed of 44 samples. We extracted the 44 Arcifera samples from Dataset E composed of 366 shared loci which present the lowest level of missingness (Table 1). First, BPP (Flouri *et al*. 2018) was used with the A11 option, using inverse-gamma distributed diffuse priors (= 3; ß = 1000) for the population sizes (θ) and root ages (τ0). Analysis was run for 100,000 generations, sampling every 100 generations after a burnin of 8,000 generations. Second, the multi-locus species delimitation using Bayesian model comparison implemented in the TR2 package (Fujisawa *et al*. 2016) has been applied on the same dataset. Locus trees generated with IQ-TREE v2.0.5 (Minh *et al*. 2020) and previously used for the weighted-ASTRAL approach were used as well as the maximum likelihood IQ-TREE consensus tree on dataset A as guide tree. Outgroups were removed from gene trees and the guide tree.

From the SNPs (Dataset H), population clustering was assessed using STRUCTURE 2.3.3 (Pritchard *et al*. 2000). Bi-allelic SNPs shared by at least 40% of the samples were extracted using VCFtools v0.1.12a (Danecek *et al*. 2011). Because markers are supposed to be unlinked, we extracted randomly only one SNP by locus. K-values from 1 to 15 were tested with no prior population information and performed three times for each of them to verify a convergence of estimations. A burn-in of 100,000 runs was used followed by 500,000 iterations. The most likely number of clusters was determined using the Evanno method (Evanno *et al*. 2005) implemented in Structure Harvester (Earl & vonHoldt 2012). The replicates were then combined and the figures generated using CLUMPAK server (Kopelman *et al*. 2015). To investigate putative admixture between species or subspecies we estimated Patterson’s D statistic (ABBA-BABA test) (Patterson *et al*. 2012) for all subspecies/species quartets using the Dsuite (Malinsky *et al*. 2021). The analyses were performed on bi-allelic SNPs shared by at least 40% of the samples composed of 6,743 SNPs. Z-scores and associated p-values were calculated to assess the significance of the results.

## Results

### Museomics approach efficiency

The combination of historical and fresh samples enabled us to compare the effectiveness of museomics methods. The DNA concentrations obtained from a single leg are very variable between fresh samples (mean = 8.37 ng/µL; sd = 6.82 ng/µL) and NHC samples (mean = 1.18 ng/µL; sd = 2.45 ng/µL). There was a significant correlation between the quantity of DNA extracted and the age of the specimens (Figure 1A). For the NHC samples, this concentration was not homogeneous, with some samples nevertheless showing a high concentration. Forty samples with a concentration below the detection thresholds were excluded from the rest of the capture process. It should be noted that some samples with very low DNA concentrations, such as *C. fabricii* CBX0094 captured in 1977 with a concentration of only 0.08 ng/µL, were reliably placed into the final phylogenetic inferences. For specimens with measurable DNA, the capture process worked efficiently, allowing the sequencing of an average of 8.4 million reads per sample (sd = 9.2 million). There was a large difference between the average number of reads obtained from fresh samples (mean = 23.8 millions; sd = 11.0 millions) and NHC samples (mean = 5.4 millions; sd = 5.0 millions). The age of the specimens also had an influence on the number of reads obtained, as there was a significant correlation between the age of the specimens and the number of reads obtained (Figure 1B).

**Figure 1.**
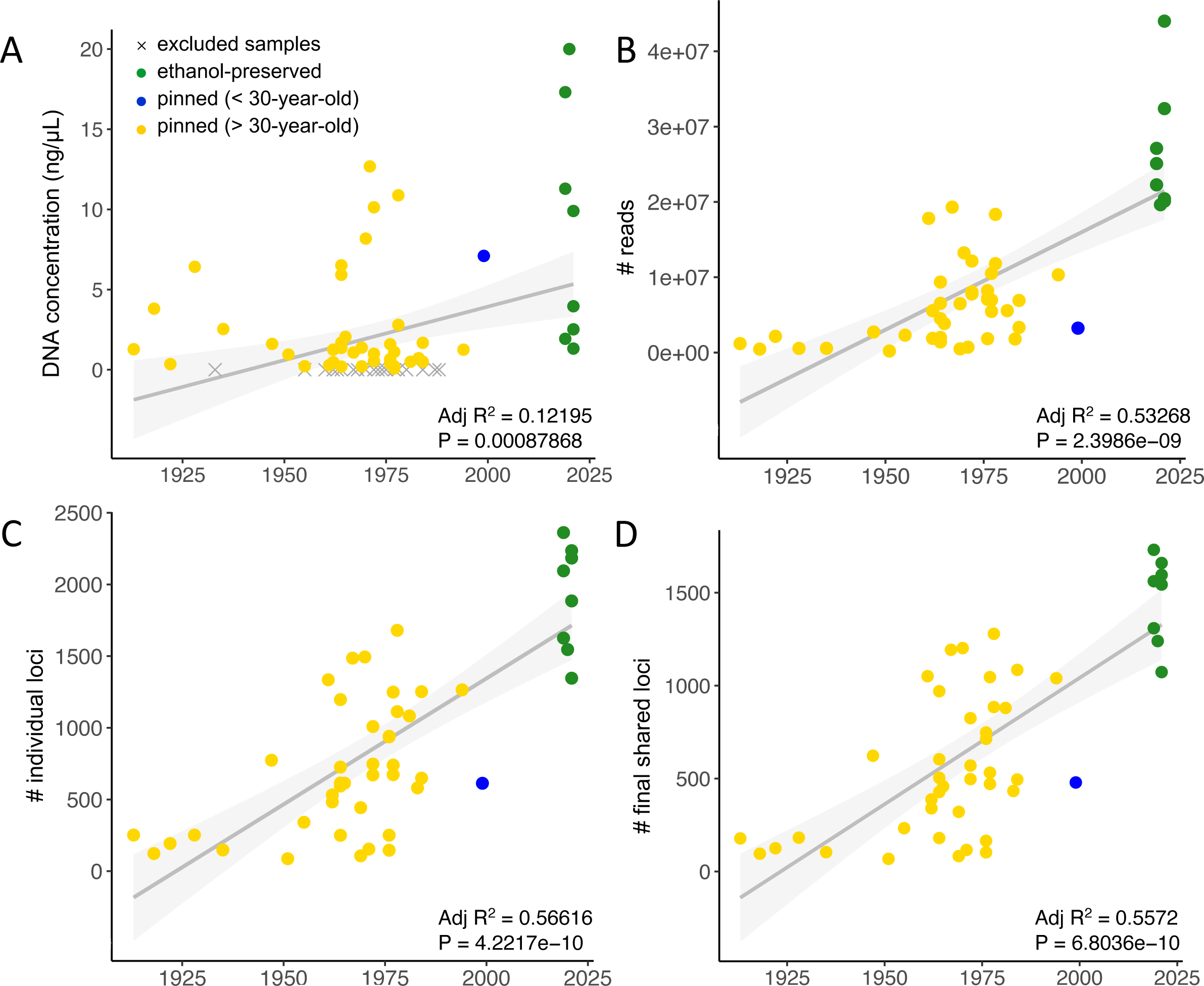
Summary of locus recovery. Plots representing the relationship between the collection year and DNA concentration (A), number of sequenced reads (B), number of loci recovered for each sample (C), and number of shared loci in final dataset (D). In each plot, ethanol-preserved samples are shown in green, samples from museums with an age < 30 years in blue and samples from museums with an age > 30 years in yellow. Correlations were tested with Spearman’s correlation tests and adjusted coefficients of determination R-squared were estimated using a linear model.

After locus reconstruction, the difference between fresh and NHC samples persists, with an average of 1765 loci recovered in fresh samples (sd = 553) and 629 in NHC samples (sd = 447) (Figure 1C). This difference is of the same order when looking at shared loci (Figure 1D). There is a large heterogeneity in the number of loci recovered between NHC samples, largely linked to the age of the specimen. Samples with too few loci (< 150 loci), i.e. 12 samples, had to be excluded from the final datasets. For 35 NHC samples, the number of loci recovered, on average 793 (sd = 400), was sufficient to include them in subsequent analyses. Although strict filtering steps reduced the number of NHC samples, they also ensured the reliability of the dataset for downstream inferences.

### Phylogenomic inferences

We inferred the phylogenetic relationships among Arcifera using six different datasets under various taxon sampling and/or gene sampling strategies (Figure 2). The results of analyses based on a concatenation approach performed in IQ-TREE and on a coalescent species-tree approach conducted in wASTRAL are consistent except for the placement of *C. marietti*, the branching pattern within *C.* (*Chaetocarabus*) and relationships between *C. creutzeri* and *C. fabricii*. The subgenus *C.* (*Hygrocarabus*) is recovered as monophyletic in all analyses (including wASTRAL) and as sister to the rest of Arcifera, however all analyses failed to recover *C. nodulosus* and *C. variolosus* as reciprocally monophyletic. In all IQ-TREE inferences except the one based on Dataset E and all wASTRAL analyses except the one based on Datasets A and B, *C. marietti* is recovered as sister to *C.* (*Chaetocarabus*) with heterogeneous levels of branch support. In the IQ-TREE analysis of Dataset E, this taxon is recovered as sister to the genus *Carabus* as a whole, whereas in wASTRAL analyses of Datasets A and B it is recovered as sister to Arcifera except *C.* (*Hygrocarabus*) with low branch support. The subgenus *C.* (*Chaetocarabus*) is always recovered as monophyletic but internal relationships differ between analyses. A minority of analyses recovered *C. arcadicus* and *C. intricatus* as reciprocally monophyletic (for instance no wASTRAL analysis recovered this relationship). The subspecies *C. intricatus lefebvrei* is recovered as sister to the nominal subspecies in all analyses. When *C. arcadicus merlini* is included (Datasets B, D and F only), it never groups with other specimens of the nominal subspecies resulting in *C. arcadicus* being consistently inferred as paraphyletic when this taxon is included (Supplementary Figure 2). The subgenus *C.* (*Platycarabus*) is recovered as monophyletic and with identical interspecific relationships across all IQ-TREE analyses but some contention in wASTRAL ones. The alpine endemic *C. cychroides* is recovered as sister to the rest of the subgenus in all analyses with strong branch support (IQ-TREE and wASTRAL). The species *C. depressus* is inferred as the next lineage branching off in *C.* (*Platycarabus*) across all IQ-TREE analyses and most wASTRAL analyses (except in Dataset A and E where it is recovered as sister to *C. irregularis* with low branch support). The subspecies *C. depressus lucens* is recovered as sister to the nominal subspecies in all analyses. The placement of the three remaining *C.* (*Platycarabus*) species is identical across all IQ-TREE analyses with strong branch support, with *C. creutzeri* being sister to *C. fabricii* and *C. irregularis*. The wASTRAL analyzes recover different relationships but with low branch support, with a weakly supported sister relationship between *C. creutzeri* and *C. fabricii* in analyses of Datasets D, E and F. The subspecies *Carabus fabricii malachiticus* is recovered as nested within the nominal subspecies in all analyses. The subspecies *Carabus irregularis montandoni* is recovered as sister to *Carabus irregularis bucephalus* and *Carabus irregularis irregularis* in all IQ-TREE analyses whereas it is *C. irregularis bucephalus* that is inferred as sister to *C. irregularis irregularis* and *C. irregularis montandoni* in all wASTRAL analyses. Overall, the IQ-TREE and wASTRAL inferences are highly compatible when collapsing the weakly supported relationships in wASTRAL species trees (gray and red asterisks in Figure 2). In particular, inconsistent relationships in wASTRAL compared to IQ-TREE always received poor branch support. We observe that branch support and overall phylogenetic resolution appears positively correlated to gene and taxon sampling (i.e., including fewer taxa and fewer loci to improve matrix completeness likely results in a loss of resolution).

**Figure 2.**
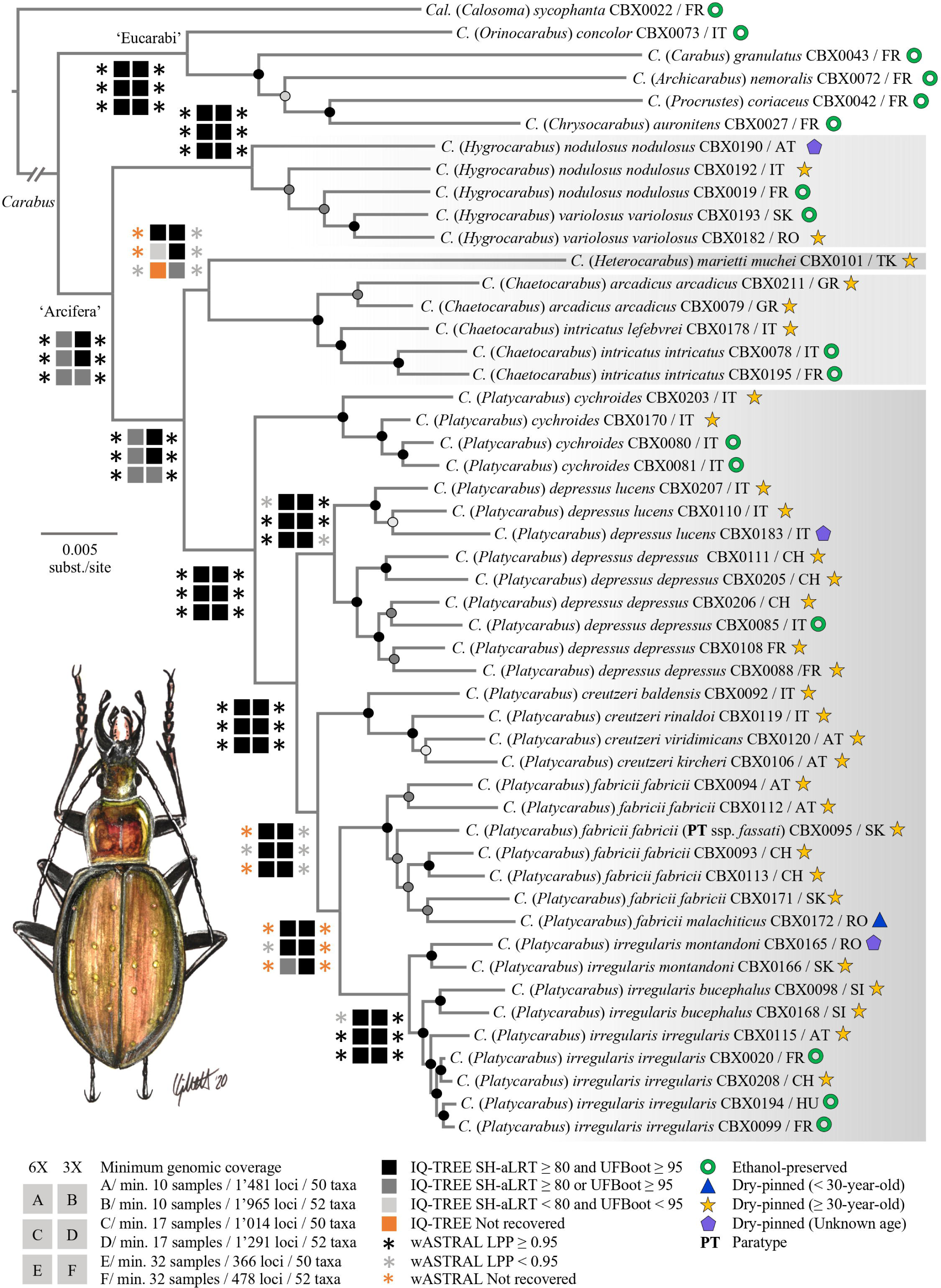
Summary of phylogenetic inferences across Arcifera based on HyRAD-X data. The presented topology is derived from a maximum likelihood analysis performed in IQ-TREE using Dataset A. Branch support from this analysis is shown for all branches. Branch support retrieved in different analyses is shown for major branches according to the inserted caption. Sample type is indicated according to the inserted caption. Abbreviations at the end of each taxon label correspond to the following countries: AT, Austria, CH, Switzerland, FR, France, GR, Greece, HU, Hungary, IT, Italy, KP, Carpathians (Slovakia to Romania), RO, Romania, SI, Slovenia, SK, Slovakia, TK, Turkey. An illustration of a male *Carabus (Platycarabus) cychroides* is presented (Drawing: Conrad Gillett).

### Divergence time estimation

The BEAST dating analysis revealed consistent results for the four main nodes, i.e. the root, *Carabus*, Arcifera and *C. (Platycarabus)* nodes, according to the three models tested, Yule, Birth-Death model, and Constant population size coalescent (Figures 3 and 4). The coalescent model including eight Bayesian log-normal relaxed clocks received the best marginal likelihood as calculated using stepping-stone sampling in BEAST and was therefore selected hereafter. This inference suggests an origin of Arcifera at 26.07 Ma (95% HPD: 22.77 - 29.67 Ma) and 14.56 Ma (95% HPD: 12.52 -16.76 Ma) for the *C. (Platycarabus)* subgenus.

**Figure 3.**
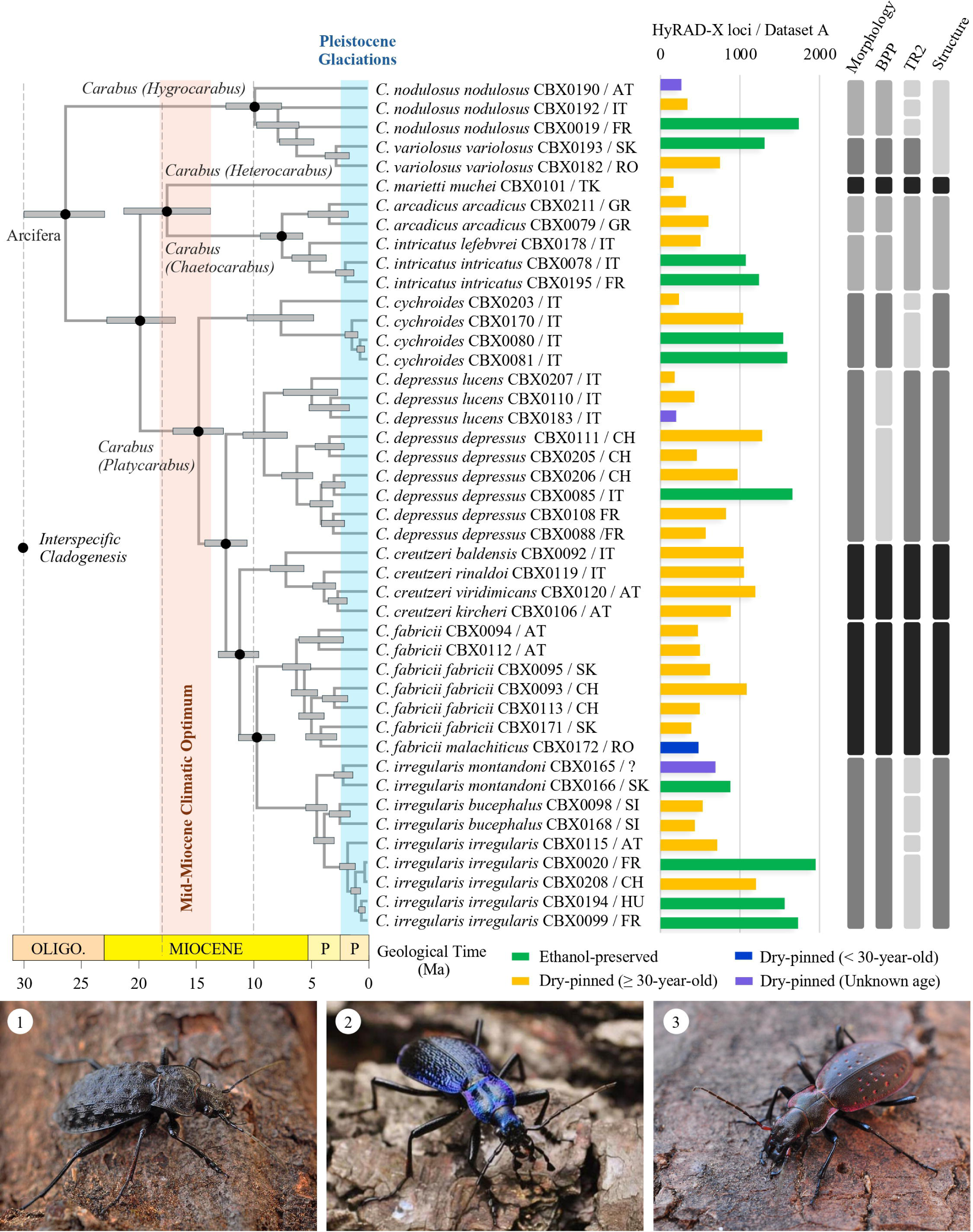
Bayesian divergence time estimates for the subgenus *Carabus (Platycarabus)* and Arcifera group. Maximum clade credibility tree obtained from a BEAST analysis using eight Bayesian log-normal relaxed clocks and a Coalescent Constant Size tree model. Node estimates are postburn in median ages, with 95% credibility intervals. Histogram represents the number of loci recovered for each sample and sample type are indicated according to the inserted caption. The section on the right shows the results of species delimitations identified using the different methods indicated above. The shades of gray represent the concordance between the different approaches with black being a total consensus. Habitus of three representative species (1) *Carabus nodulosus nodulosus* (credit: Conrad Gillett), (2) *Carabus intricatus intricatus* (credit: Conrad Gillett) and (3) *Carabus irregularis irregularis* (credit: Conrad Gillett) are shown.

**Figure 4.**
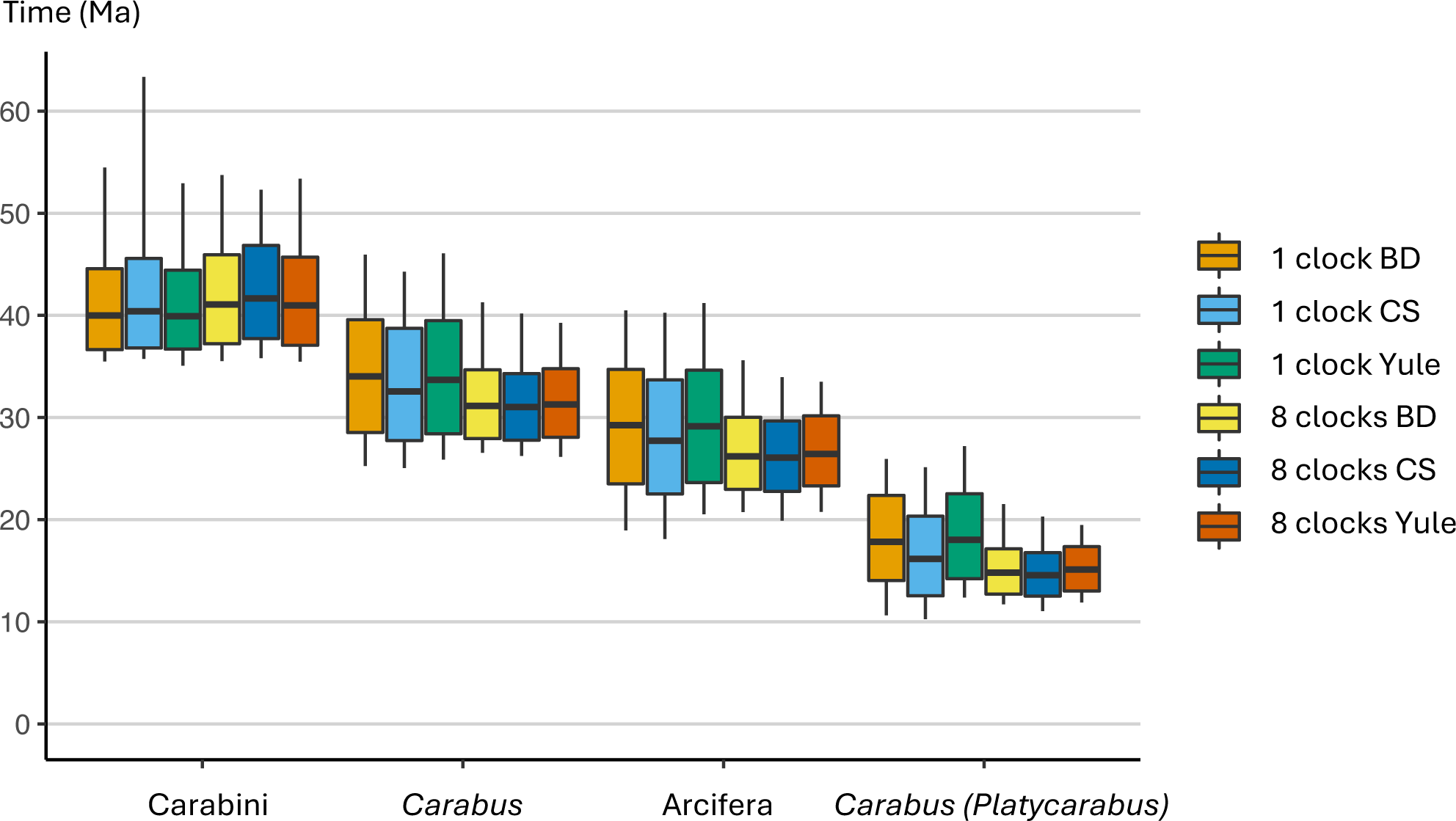
Comparison of divergence time estimation between competing tree models and relaxed-clock partitioning strategies. Box-plots indicate for each analysis (color-coding inserted as a caption on the right side of the figure) the median age of the focal node (see X axis) and associated 95% age credibility interval. BD, birth-death model; CS, constant population size coalescent model.

### Species delimitation and putative hybridization

The different approaches to species delimitation produced contrasting results. The analysis performed with BPP is the most consistent with morphology and the current classification. The two species *C. variolosus* and *C. nodulosus* are well separated even though *C. nodulosus* is not monophyletic in our phylogeny. TR2 approach proposes an oversplit of the three *C. nodulosus* samples. Conversely, the STRUCTURE approach groups the two species in a single cluster (Supplementary Figure 6). STRUCTURE analyses reveal that the most probable number of clusters is found for K=5 (deltaK = 1539.43) (Supplementary Figure 6). However, some clear splits emerge at higher K values. This is the case for *C. (Heterocarabus) marietti*, represented by a single sample which is identified as a separate cluster at K=7 or the split between *C. (Chaetocarabus) arcadicus* -*C. (Chaetocarabus) intricatus* and *C. (Platycarabus) cychroides* which emerges at K values greater than 5. The divisions we propose (Figure 3) are therefore a combination of the logical splits identified by STRUCTURE for the different values of K. *C. marietti*, the only representative of *C. (Heterocarabus)* is delineated as a species in all three approaches. Within *C. (Chaetocarabus)*, the two species *C. arcadicus* and *C. intricatus* are delineated by BPP but are merged by TR2 and STRUCTURE, potentially for the same reasons as in *C. (Hygrocarabus)*. It should be noted that the two subspecies of *C. intricatus*, i.e. *C. intricatus lefebvrei* and *C. intricatus intricatus*, are never delineated as distinct species. The species *C. cychroides* was well discriminated in two of the three approaches, with only TR2 proposing an additional split of the most basal sample. The two subspecies of *C. depressus*, *C. depressus depressus* and *C. depressus lucens* are not grouped together in the BPP approach and are identified as two distinct species. The results of the three methods are fully consistent with the morphology for *C. creutzeri* and *C. fabricii*. For *C. irregularis*, the situation is similar for two of the three methods, BPP and Structure. Among the 85 trios analyzed, high D-statistics values, > 0.25, with significant p-values were observed for three trios. For two of these, *C. cychroides* was observed in P1 and *C. arcadicus* in P3. Despite this, no f-branch signal significantly different from zero could be identified (Supplementary Figure 7). These results suggest an absence of past introgression between the different species and subspecies.

## Discussion

### Using museomics to obtain an extensive dataset

The HyRAD and HyRAD-X methods are unique in that they allow in-house production of probes using a ddRAD protocol, either directly on the DNA of a few fresh samples (Suchan *et al*. 2016) or on their RNA (Schmid *et al*. 2017). These approaches allow targeting of several thousand loci and, in turn, generation of high-resolution phylogenomic inferences (Young & Gillung 2020). In this study, we obtained 1,965 loci for the most extensive dataset. These loci were informative enough to resolve both the deep relationships between outgroups and the more recent relationships at the intrageneric and intraspecific scales. In addition, the identification of SNPs on these loci also enabled population genomic approaches such as the study of genetic structure and admixture.

In addition, the HyRAD-X approach made it possible to integrate samples with extremely low initial DNA quantities. However, out of 96 samples from which DNA was extracted, 40 had an undetectable quantity of DNA. In the context of museomics projects, it is therefore instrumental to plan for redundancy in the sampling, with several samples per targeted taxon, in order to compensate for any failures. Furthermore, the ability to generate genetic information from hDNA is not entirely predictable. The recovery of meaningful genomic data does not seem to be linked to the age of samples (Figure 1), in line with existing observations (Toussaint *et al*. 2021; Nunes *et al*. 2022). In that vein, large amounts of genomic data could be obtained from some older NHC samples when almost none could be obtained from more recent samples. The quality and quantity of DNA that can be extracted from NHC specimens is linked to factors that we cannot control, such as the conditions of collection and preservation process (Post *et al*. 1993; Dillon *et al*. 1996; Ruppert *et al*. 2023).

### Systematics and species delimitation in Arcifera

Our results provide a robust phylogenomic tree of Arcifera for the first time (Figure 2). Overall, we support the view that Arcifera represents a monophylum within which all four subgenera form clades. The monophyly of Arcifera is also supported by the presence of a hook-shaped ligulum (i.e., arculus) at the base of the endophallus, a strong morphological character that unites all constituents of this lineage (Imura *et al*. 2000; Deuve *et al*. 2012). Our study is the first to provide strong evidence for these relationships while including all species of the group. Other studies based on reduced genomic sampling, often a single gene fragment, either failed to recover Arcifera as monophyletic (Imura *et al*. 2000; Osawa *et al*. 2004), or had too limited a taxon sampling to properly test the placement and otherwise monophyly of each subgenus (Su *et al*. 2003; Sota & Ishikawa 2004; Deuve *et al*. 2012). Except for a minority of analyses, our results strongly suggest that *C. (Hygrocarabus*) is sister to the rest of Arcifera, with *C. (Platycarabus)* as sister to a clade formed by *C*. (*Chaetocarabus*) and *C. (Heterocarabus)*.

Within *C. (Hygrocarabus*), we recover *C. variolosus* nested within *C. nodulosus*. This result contrasts with Mossakowski *et al*. (2020) where the two species were suggested to be well differentiated genetically. In their study, these authors argued based on the analysis of two gene fragments that both taxa form distinct clades although several specimens caused each species to be paraphyletic. Some tests of mating between the two candidate species were also performed in this study and suggested that the two lineages do not mate. However, the scale and conditions of these trials do not allow to conclusively rule out potential mating. We argue that in the current state of our knowledge it is not yet possible to definitively test species boundaries, past introgression and signature of hybridization between *C. nodulosus* and *C. variolosus*. A desired approach would be to combine large geographical sampling as in Mossakowski *et al*. (2020) with a genomic scale dataset as developed in the present study to revisit the systematic conundrum within this subgenus at the population level.

Within *C.* (*Chaetocarabus*), we recover *C. arcadicus* as sister to *C. intricatus* in most analyses (Figure 2). These two species are allopatric, morphologically well-differentiated and little doubt exists with respect to their status as distinct species. Surprisingly our species delimitation analyses only partly support the two species hypothesis, with TR2 and STRUCTURE considering that *C. (Chaetocarabus*) is a unique species. Considering the low genomic coverage of some taxa included in the analyses (see below), the clear morphological and geographical split between these lineages and the support from BPP analyses, we argue that the validity of these two species is uncontroversial. Natural hybrids with an intermediate morphology and usually green dorsal pattern are known to exist along the limits of their respective ranges in northern Greece (i.e., at the Katara pass) where *Carabus intricatus macedonicus* (not sampled here) and *C. arcadicus arcadicus* co-occur. Both *C. arcadicus* and *C. intricatus* also comprise geographically restricted subspecies in Greece that have been considered valid species by some authors (e.g., Ishikawa 1984; Baviera & Micali 2021). In the south of Greece, the melanistic subspecies *C. arcadicus merlini* is endemic to the Peloponnese peninsula and allopatric from the nominal subspecies present in the north (Arndt *et al*. 2011). One specimen of this taxon was sequenced but genomic coverage was low and therefore it was only included in the less stringent Datasets B, D and F. In the phylogenetic analyses of these datasets, the inclusion of *C. arcadicus merlini* systematically results in all three *C. arcadicus* specimens forming a phylogenetic grade within which *C. intricatus* is nested. We argue that this is an artifact possibly caused by missing genomic sampling and that both species are reciprocally monophyletic as recovered in all other analyses and as suggested by morphology. However, it is possible that *C. arcadicus merlini* represents a distinct evolutionary lineage since it is always recovered as sister to the rest of *C. (Chaetocarabus)*. Additional taxon sampling is needed to test the placement of this morphologically distinct taxon within the subgenus.

Across its range, *C. intricatus* is represented by the nominal subspecies from western France and UK to northern Greece. In the south of Italy and Sicily, this species is represented by the allopatric *C. intricatus lefebvrei*. The status of this taxon is debated, and some authors consider it a valid species (e.g., Ishikawa 1984; Giglio *et al*. 2013; Baviera & Micali 2021; Talarico et al. 2021). In our results, we recover this subspecies as sister to the nominal subspecies represented by specimens from France and Piemonte. Our phylogenetic inferences support the view of *C. intricatus lefebvrei* as a possible distinct species but our species delimitation analyses reject this hypothesis. To properly test species boundaries within *C. intricatus*, additional taxon sampling is needed including a much denser geographical sampling of the nominal subspecies along with all described valid subspecies (Deuve 2019). In the Balkans, several subspecies of *C. intricatus* have been described and represented more or less isolated populations restricted to northern Greece. Despite our efforts we could not obtain DNA of good quality for *C. intricatus krueperi* endemic to eastern Thessaly and considered by some authors to be a valid species. Here as well, a denser taxon sampling is needed to properly test species boundaries in this group.

The placement of *C. (Heterocarabus) marietti* as sister to *C.* (*Chaetocarabus*) receives support from most analyses in this study. Despite a relatively circumscribed geographic range in northern Turkey and southern Bulgaria, numerous taxa have been described in this subgenus even though currently a single species is considered valid (Turin *et al*. 2003; Deuve 2019). Increasing the taxon sampling for this group by covering all its geographical range would allow testing the match between morphological and genetic diversity and better understand the evolution of this unique lineage at the inter- and intraspecific interface.

Within *C. (Platycarabus)*, we recover *C. cychroides* as sister to the rest of the subgenus. This result is unexpected because this species is a very narrowly restricted endemic to Piemonte mountain ranges where it lives in alpine meadows and scree >2000m. The species was only included once in a phylogenetic framework by Casale *et al*. (1998) who recovered it as a derived lineage close to *C. depressus* and *C. fabricii*. Interestingly, a sister relationship of this species to the rest of *C. (Platycarabus)* was suggested by the analysis of morphological characters in Casale *et al*. (1998). Indeed, this species is morphologically quite different from the rest of the subgenus in that it is one of the most extreme examples of cychrization in *Carabus*. All species of the subgenus present a stenocephalic morphology, although less marked than in *C. cychroides*, except for *C. irregularis* which is macrocephalic. Our phylogenetic inferences are therefore important to understanding the evolution of predation strategies and associated morphology across the genus *Carabus* in which both types of morphologies exist (Sota & Ishikawa 2004). Most malacophagous and helicophagous species in *Carabus* are macrocephalic and use their enlarged pronotum, head and robust mandibles to break snail shells. Cases of stenocephaly are most notably observed in *C. (Platycarabus)* but also in *Carabus* (*Damaster*) Kollar, 1836 and *Carabus* (*Macrothorax*) Desmarest, 1850. The fact that *C. irregularis*, the only *C.* (*Platycarabus*) macrocephalic species, is recovered as the most derived species in the subgenus, indicates that macrocephaly possibly evolved from a stenocephalic morphology unlike what was suggested in Casale *et al*. (1998). In the case of *C. cychroides*, it is not closely related to any other species of the subgenus as suggested by previous authors, and despite rare known natural hybrids with *C. depressus* in the Cottian Alps (i.e., Colle delle Finestre, Monte Morefreddo, Monte Albergian), these species do not share an immediate recent common ancestry (Sturani 1962; Casale *et al*. 1998; Anselmo & Rizzioli 2022a; b).

The rest of *C. (Platycarabus)* species and most sampled subspecies are found to be monophyletic (Figure 2). We recover the subspecies *C. depressus lucens* as sister to the nominal subspecies in all analyses and with robust branch support. This subspecies is morphologically quite divergent from the nominal subspecies and *C. depressus bonellii* as it completely lacks elytral foveae. It is also allopatric from the rest of the *C. depressus* populations, being found in a small transalpine region between France and Italy (i.e., French Queyras to Italian Alpi Marittime), and its status as a valid species, even though rejected by three out of four species delimitation analyses, should be revisited with enhanced population sampling. Our taxon sampling within *C. creutzeri* does not allow testing subspecies monophyly and relationships in detail but species delimitation analyses unambiguously support a single species (Figure 3). Within *C. fabricii*, we recover the Carpathian populations of *C. fabricii* (ssp *fassati* = nominal ssp, and spp *malachiticus*) nested within Alpine populations of the nominal subspecies. This is unexpected to some extent as *C. fabricii* presents a disjunct distribution between the Alps and the Carpathians (i.e., it is not currently found in the Danube valley). Our results suggest that despite an allopatric range, gene flow has been maintained between all populations of this species, however branch supports for internal relationships in *C. fabricii* are moderate and enhanced taxon sampling is needed to understand the past and present connectivity between populations. All species delimitation analyses support a unique species. One of the most interesting subspecific cases is recovered in *C. irregularis*. This species is the most widespread of the subgenus ranging from eastern France to Romania and Ukraine. It comprises three valid subspecies, one of which *C. irregularis montandoni* from the Carpathians, was suggested to be a valid species based on molecular evidence (Homburg *et al*. 2013). Our results support to some extent this view, with *C. irregularis montandoni* being found sister to the rest of populations in all IQ-TREE analyses but not in wASTRAL analyses where the other subspecies *C. irregularis bucephalus* is found as sister to the rest of the clade. There seems to be a genetic differentiation between the three recognized subspecies of *C. irregularis* but our species delimitation analyses support the view of a single species.

### Evolution of the Arcifera group

The divergence time estimation analyses all recover an origin of Arcifera ca. 26 Ma in the Oligocene. We did not perform a biogeographic estimation of ancestral ranges in the group because several species are very widespread and initial attempts resulted in unresolved patterns. The fact that species boundaries within *C. (Hygrocarabus*) are unstable also prevented a proper reconstruction. However, it is possible to discuss several phylogenetic splits in the framework of our results. The stem branch connecting *C. (Hygrocarabus*) to the rest of Arcifera is long, potentially representing periods of extinction in this lineage. Currently the two recognized species in the subgenus occur in temperate forests where adults live and hunt near and in good quality streams. The reconfiguration of such habitats in the past 25 million years due to climatic oscillations (Westerhold *et al*. 2020) may have extirpated populations and pushed others into their current ranges. Considering the specificity of these two lineages to their habitat, and predictions of global warming and their impact on such ecosystems (Capon *et al*. 2021; Bonacina *et al*. 2023), it is likely that they may be increasingly threatened in the future.

With respect to biogeography, one of the most interesting lineages in Arcifera is the clade composed of *C. (Chaetocarabus*) and *C. (Heterocarabus)*. Because *C. (Heterocarabus) marietti* is restricted to eastern Bulgaria and western Turkey, and *C. (Chaetocarabus*) distributed in Greece (*C. arcadicus* is endemic to Greece), it is likely that the ancestors of this clade originated in the geologically highly complex Aegean area. The split between the two subgenera ca. 17 Ma predates the timing of the opening of the Aegean Sea in the Tortonian ca. 8 Ma (i.e., opening of the Mid-Aegean Trench or Aegean barrier; van Hinsbergen & Schmid 2012), rejecting the hypothesis of geographic vicariance in the south as suggested in other lineages (Poulakakis *et al*. 2015). Interestingly, both subgenera have very marginally overlapping distributions in the Thrace basin with *C. (Heterocarabus)* currently distributed on the southern Black Sea coast where *Carabus intricatus* is also represented by the subspecies *C. intricatus starensis* (Gueorguiev & Gueorguiev 1995). At the time of divergence in the early Miocene (i.e., Burdigalian), the Thrace basin formed a connection between the eastern Balkan peninsula and Anatolia (Rögl 1997, 1999; Sachsenhofer *et al*. 2017; Erbil *et al*. 2021). It is possible that ancestral populations dispersed in the Balkan Peninsula and/or in Anatolia where they evolved independently. Under this scenario, the close geographic ranges of these two species would likely represent secondary contact associated with more recent colonization of the Thrace basin. More robust population-level taxon sampling, especially of *C. (Heterocarabus)*, might elucidate the fine-scale biogeographic history of this clade in the future. Within *C. (Chaetocarabus*), the two currently recognized species are mostly allopatric with only a short overlap in western Greece (e.g. Katara pass). There is no clear geological barrier that may have fostered vicariant diversification at the time of speciation ca. 7 Ma. Further diversification appears to be occurring at the population level with *C. intricatus lefebvrei* endemic to southern Italy and allopatric from the nominal subspecies. Similarly, *C. arcadicus merlini* endemic to Peloponnese is morphologically quite divergent from the nominal subspecies and might represent a case of ongoing speciation. The wide dispersal of *C. intricatus* across the western Palearctic region is likely recent and may be explained by the generalist habitat preference of this species. Additional geographical and taxon sampling will likely yield more robust inferences of evolutionary patterns and processes within this clade in the future.

The evolutionary history of the subgenus *C. (Platycarabus)* is also revealed by our analyses. We recover the narrowly endemic *C. cychroides* as sister to the rest of the subgenus. This is surprising as it was not suggested by the molecular inference of Casale *et al*. (1998). This placement has strong implications for our understanding of alpine biogeography in this group. Only *C. irregularis* has lowland populations and its derived placement in the phylogeny indicates that alpine specialization was likely ancestral in the subgenus with recent shift in that species to lower habitats. This phylogenomic pattern and the origin of the subgenus *ca.* 15 Ma during the warmest period of the Neogene seems to indicate that ancestors of *C. (Platycarabus)* may have been less specialized than nowadays and were distributed in mountain regions. In the mid Miocene, mountain ranges across the Alps had the same elevation as nowadays (Campani *et al*. 2012; Krsnik *et al*. 2021), however ecosystems were different due to significantly warmer climatic conditions. When the climate progressively turned colder these beetles adapted to ensuing conditions and became alpine specialists. It is possible that species of the subgenus diverged due to competition, niche filling and/or host specialization as observed in *C. cychroides* for instance. We hypothesize that in the latest sequence of their evolutionary history, Pleistocene glaciations played a limited role in speciation since all current species had already diverged (Figure 3).

Although natural hybrids are known between different species of the subgenus, our results recover no hybridization signal between them. The most significant case concerns the species *C. fabricii* and *C. irregularis*, whose ranges largely overlap in Switzerland, Austria and Slovakia. It is in these sympatric areas that several cases of natural hybridization have been identified (e.g. at the Radstädter Tauern Pass in Austria, Mandl 1960). However, our genetic results do not show any hybridization signals between the species, either on genetic structure, where the two clusters are well separated, or in the approach using Dsuite, which seeks to trace admixture signals in the lineages. These results suggest that these sporadic hybridization events are not conserved in populations and could imply a potential infertility of F1s (Casale *et al*. 1998). Furthermore, the networks obtained with the three mitochondrial genes (Supplementary Figure 4) group the samples of the *C. irregularis* and *C. fabricii* species in the same cluster. These mitonuclear discordance patterns are frequent in the literature and can be explained by the specific biological properties of mitochondrial DNA (uniparental inheritance and reduced recombination; Birky 2001) or differences in the evolutionary histories of nuclear and mitochondrial markers including incomplete lineage sorting and gene flow among species (Sota & Vogler 2001; Suchan *et al*. 2017). The results obtained with the nuclear loci are sufficiently robust to be able to consider that the cases of hybridization observed are either localized or do not induce lasting admixture between the species. A more detailed analysis of hybrids, local populations and the implications of hybridization on the fitness of individuals could provide a better understanding of the mechanisms involved.

Integrating current species distribution, genetic isolation of these alpine species was already in place when glaciation cycles struck the Alps. As a result, dispersal of populations in peripheral glacial refugia as observed in *C. irregularis* (Homburg *et al*. 2013) did not result in genetic homogenization despite species being placed in secondary contact. It is also possible in the case of the more alpine-adapted species (all but *C. irregularis*) that dispersal occurred in nunataks rather than peripheral glacial refugia (Holderegger & Thiel-Egenter 2009; Schönswetter & Schneeweiss 2019; Kosiński *et al*. 2019), which would have resulted in increased genetic differentiation among populations as suggested by our analyses. Coupling more extensive geographic sampling of these five alpine species with niche modeling analyses may help testing more specifically the different scenarios that governed range and genetic evolution of these populations during Pleistocene glaciations.

## Acknowledgements

We warmly thank Michael Balke for the loan of material from the ZSM-SNSB. We thank Elsa Ricossa for the digitization of specimens housed at the Natural History Museum of Geneva. We thank Céline Rochet for assistance in fieldwork. We thank the city of Geneva for an internal student grant awarded to MTP. We thank Conrad Gillett for allowing the use of his photographs and drawings in this article. We thank Ivan Rapuzzi for fruitful discussions and feedback on taxonomic aspects of this work. We also thank Michael Caterino, Julian Dupuis and an anonymous reviewer for insightful comments on an earlier version of this article, as well as Felix Sperling for his editorial work.

## Data, script, code, and supplementary information availability

Raw reads are available on the NCBI BioProject PRJNA1086379. The data underlying this article (final alignments and trees) and bioinformatic scripts are available on Github repository (https://github.com/JeremyLGauthier/Arcifera_phylogeny).

## Figure captions

**Supplementary Table 1.** Descriptive statistics for each included and non-included sample, including historical sample data, molecular biology information (DNA concentrations), sequencing and loci reconstruction statistics.

**Supplementary Figure 1.** Schematic representation of the bioinformatic pipeline.

**Supplementary Figure 2.** Maximum likelihood trees for each dataset: Dataset A (min_cov=6, min_sample=10, 50 taxa, 1’481 loci), Dataset B (min_cov=6, min_sample=10, 52 taxa, 1’965 loci), Dataset C (min_cov=6, min_sample=17, 50 taxa, 1’014 loci), Dataset D (min_cov=3, min_sample=17, 52 taxa, 1’291 loci), Dataset E (min_cov=6, min_sample=32, 50 taxa, 366 loci) and Dataset F (min_cov=3, min_sample=32, 52 taxa, 478 loci). Node supports indicate SH-aLRT and UFBoot values.

**Supplementary Figure 3.** Species trees obtained with wASTRAL on each dataset: Dataset A (min_cov=6, min_sample=10, 50 taxa, 1’481 loci), Dataset B (min_cov=6, min_sample=10, 52 taxa, 1’965 loci), Dataset C (min_cov=6, min_sample=17, 50 taxa, 1’014 loci), Dataset D (min_cov=3, min_sample=17, 52 taxa, 1’291 loci), Dataset E (min_cov=6, min_sample=32, 50 taxa, 366 loci) and Dataset F (min_cov=3, min_sample=32, 52 taxa, 478 loci). Node supports indicate SH-aLRT and UFBoot values. Node supports indicate LPP values.

**Supplementary Figure 4.** Individual locus haplotype networks (A. CO1, B. CO3 and C. CYTB). Networks were generated in SplitsTree using calculated uncorrected p-distances and the NeighborNet algorithm. The color ellipses delimit the morphological species groups (photo credit: Marie Pauli).

**Supplementary Figure 5.** Species trees obtained with RAxML on Dataset H (min_cov=3, min_sample=4, 44 taxa, 26,201 SNPs). Node supports indicate Bootstrap values.

**Supplementary Figure 6.** Structure plots estimated on unlinked shared SNPs for K=1 to K = 15. For each K, the Mean(LnProb) is indicated.

**Supplementary Figure 7.** F4-branch statistic plotted as a heatmap. The tree topology is plotted above, and on the left, every branch of the tree is displayed (including internal branches).

