## Supplementary Figures for "Museomics of *Carabus* giant ground beetles shows an Oligocene origin and *in situ* alpine diversification"

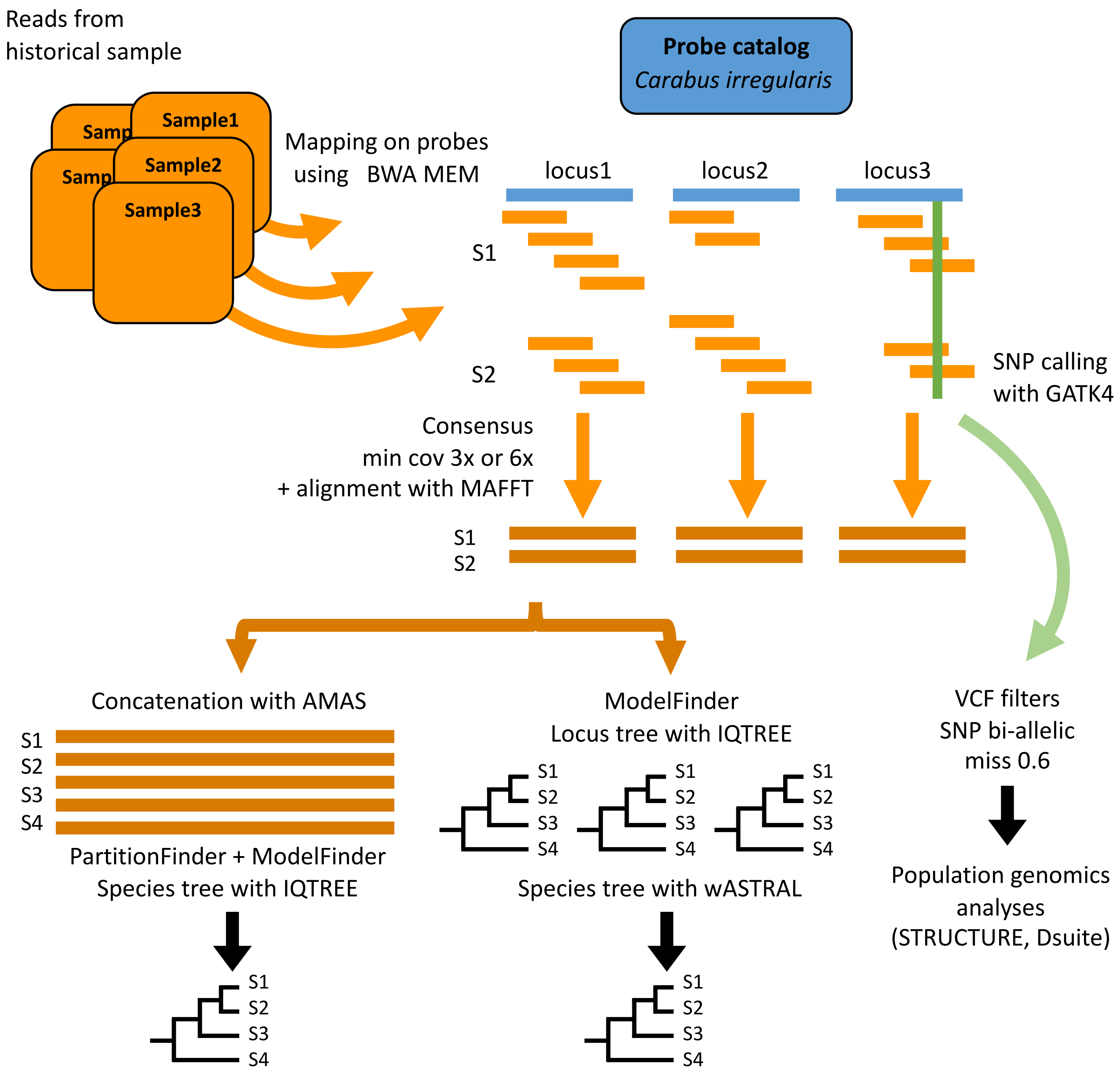


Supplementary Figure 1.

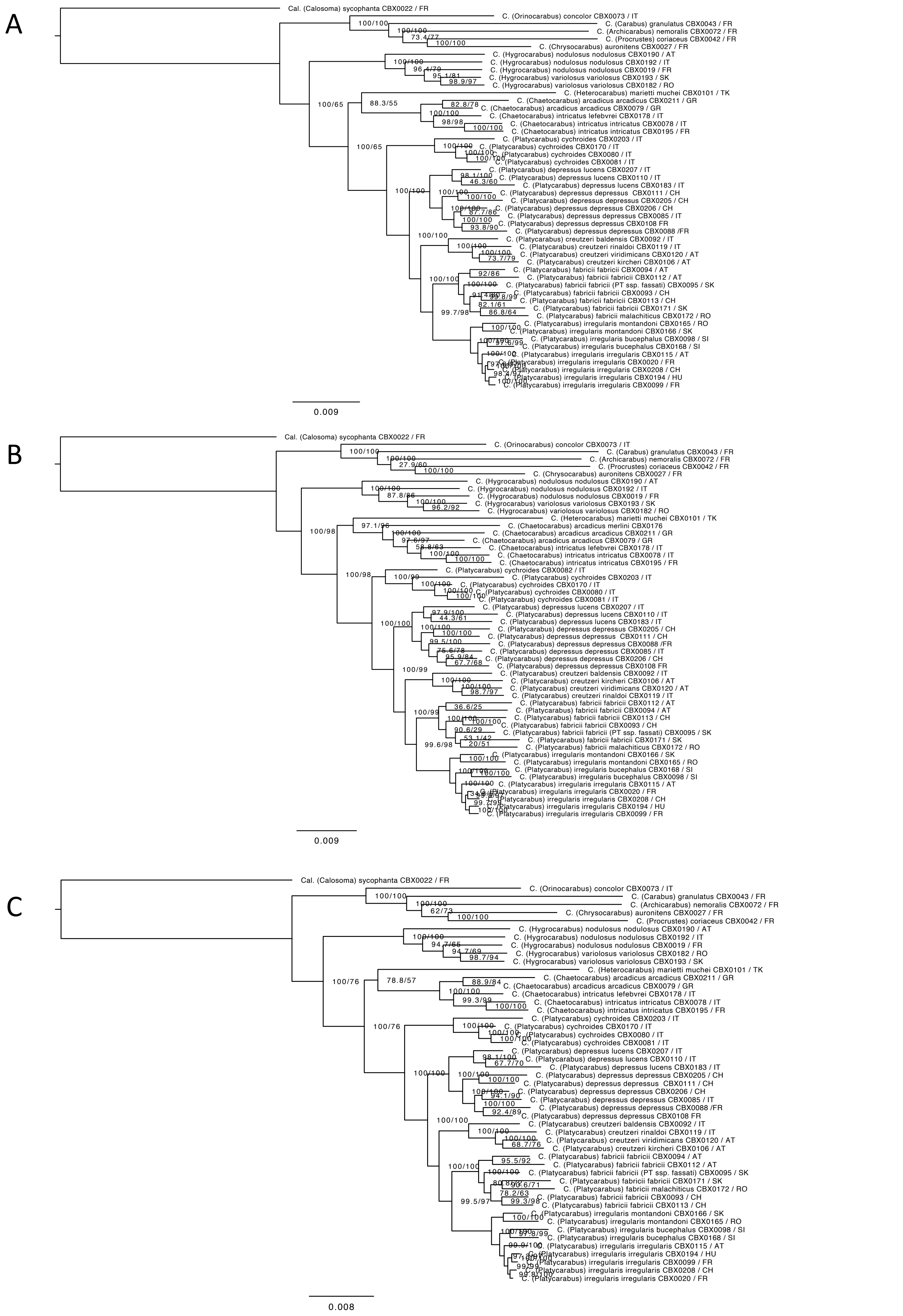
A
Cal. (Calosoma) sycophanta CBX0022 / FR
C. (Orinocarabus) concolor CBX0073 / IT
C. (Carabus) granulatus CBX0043 / FR
C. (Archicarabus) nemoralis CBX0072 / FR
C. (Procrustes) coriaceus CBX0042 / FR
C. (Chrysocarabus) auronitens CBX0027 / FR
C. (Hygrocarabus) nodulosus nodulosus CBX0190 / AT
C. (Hygrocarabus) nodulosus nodulosus CBX0192 / IT
C. (Hygrocarabus) nodulosus nodulosus CBX0019 / FR
C. (Hygrocarabus) variolosus variolosus CBX0193 / SK
C. (Hygrocarabus) variolosus variolosus CBX0182 / RO
C. (Heterocarabus) marietti muchei CBX0101 / TK
C. (Chaetocarabus) arcadicus arcadicus CBX0211 / GR
C. (Chaetocarabus) arcadicus arcadicus CBX0079 / GR
C. (Chaetocarabus) intricatus lefebvrei CBX0178 / IT
C. (Chaetocarabus) intricatus intricatus CBX0078 / IT
C. (Chaetocarabus) intricatus intricatus CBX0195 / FR
C. (Platycarabus) cychroides CBX0203 / IT
C. (Platycarabus) cychroides CBX0170 / IT
C. (Platycarabus) cychroides CBX0080 / IT
C. (Platycarabus) cychroides CBX0081 / IT
C. (Platycarabus) depressus lucens CBX0207 / IT
C. (Platycarabus) depressus lucens CBX0110 / IT
C. (Platycarabus) depressus lucens CBX0183 / IT
C. (Platycarabus) depressus depressus CBX0111 / CH
C. (Platycarabus) depressus depressus CBX0205 / CH
C. (Platycarabus) depressus depressus CBX0206 / CH
C. (Platycarabus) depressus depressus CBX0085 / IT
C. (Platycarabus) depressus depressus CBX0108 FR
C. (Platycarabus) depressus depressus CBX0088 /FR
C. (Platycarabus) creutzeri baldensis CBX0092 / IT
C. (Platycarabus) creutzeri rinaldoi CBX0119 / IT
C. (Platycarabus) creutzeri viridimicans CBX0120 / AT
C. (Platycarabus) creutzeri kircheri CBX0106 / AT
C. (Platycarabus) fabricii fabricii CBX0094 / AT
C. (Platycarabus) fabricii fabricii CBX0112 / AT
C. (Platycarabus) fabricii fabricii (PT ssp. fassati) CBX0095 / SK
C. (Platycarabus) fabricii fabricii CBX0093 / CH
C. (Platycarabus) fabricii fabricii CBX0113 / CH
C. (Platycarabus) fabricii fabricii CBX0171 / SK
C. (Platycarabus) fabricii malachiticus CBX0172 / RO
C. (Platycarabus) irregularis montandoni CBX0165 / RO
C. (Platycarabus) irregularis montandoni CBX0166 / SK
C. (Platycarabus) irregularis bucephalus CBX0098 / SI
C. (Platycarabus) irregularis bucephalus CBX0168 / SI
C. (Platycarabus) irregularis irregularis CBX0115 / AT
C. (Platycarabus) irregularis irregularis CBX0020 / FR
C. (Platycarabus) irregularis irregularis CBX0208 / CH
C. (Platycarabus) irregularis irregularis CBX0194 / HU
C. (Platycarabus) irregularis irregularis CBX0099 / FR
100/100
73.4/77
100/65
88.3/55
100/65
100/100
99.7/98
0.009
B
Cal. (Calosoma) sycophanta CBX0022 / FR
C. (Orinocarabus) concolor CBX0073 / IT
C. (Carabus) granulatus CBX0043 / FR
C. (Archicarabus) nemoralis CBX0072 / FR
C. (Procrustes) coriaceus CBX0042 / FR
C. (Chrysocarabus) auronitens CBX0027 / FR
C. (Hygrocarabus) nodulosus nodulosus CBX0190 / AT
C. (Hygrocarabus) nodulosus nodulosus CBX0192 / IT
C. (Hygrocarabus) nodulosus nodulosus CBX0019 / FR
C. (Hygrocarabus) variolosus variolosus CBX0193 / SK
C. (Hygrocarabus) variolosus variolosus CBX0182 / RO
C. (Heterocarabus) marietti muchei CBX0101 / TK
C. (Chaetocarabus) arcadicus merlini CBX0176
C. (Chaetocarabus) arcadicus arcadicus CBX0211 / GR
C. (Chaetocarabus) arcadicus arcadicus CBX0079 / GR
C. (Chaetocarabus) intricatus lefebvrei CBX0178 / IT
C. (Chaetocarabus) intricatus intricatus CBX0078 / IT
C. (Chaetocarabus) intricatus intricatus CBX0195 / FR
C. (Platycarabus) cychroides CBX0082 / IT
C. (Platycarabus) cychroides CBX0203 / IT
C. (Platycarabus) cychroides CBX0170 / IT
C. (Platycarabus) cychroides CBX0080 / IT
C. (Platycarabus) cychroides CBX0081 / IT
C. (Platycarabus) depressus lucens CBX0207 / IT
C. (Platycarabus) depressus lucens CBX0110 / IT
C. (Platycarabus) depressus lucens CBX0183 / IT
C. (Platycarabus) depressus depressus CBX0205 / CH
C. (Platycarabus) depressus depressus CBX0111 / CH
C. (Platycarabus) depressus depressus CBX0088 /FR
C. (Platycarabus) depressus depressus CBX0085 / IT
C. (Platycarabus) depressus depressus CBX0206 / CH
C. (Platycarabus) depressus depressus CBX0108 FR
C. (Platycarabus) creutzeri baldensis CBX0092 / IT
C. (Platycarabus) creutzeri kircheri CBX0106 / AT
C. (Platycarabus) creutzeri viridimicans CBX0120 / AT
C. (Platycarabus) creutzeri rinaldoi CBX0119 / IT
C. (Platycarabus) fabricii fabricii CBX0112 / AT
C. (Platycarabus) fabricii fabricii CBX0094 / AT
C. (Platycarabus) fabricii fabricii CBX0113 / CH
C. (Platycarabus) fabricii fabricii CBX0093 / CH
C. (Platycarabus) fabricii fabricii (PT ssp. fassati) CBX0095 / SK
C. (Platycarabus) fabricii fabricii CBX0171 / SK
C. (Platycarabus) fabricii malachiticus CBX0172 / RO
C. (Platycarabus) irregularis montandoni CBX0166 / SK
C. (Platycarabus) irregularis montandoni CBX0165 / RO
C. (Platycarabus) irregularis bucephalus CBX0168 / SI
C. (Platycarabus) irregularis bucephalus CBX0098 / SI
C. (Platycarabus) irregularis irregularis CBX0115 / AT
C. (Platycarabus) irregularis irregularis CBX0020 / FR
C. (Platycarabus) irregularis irregularis CBX0208 / CH
C. (Platycarabus) irregularis irregularis CBX0194 / HU
C. (Platycarabus) irregularis irregularis CBX0099 / FR
100/100
27.9/60
100/98
97.1/96
100/98
100/99
100/100
100/99
99.6/98
0.009
C
Cal. (Calosoma) sycophanta CBX0022 / FR
C. (Orinocarabus) concolor CBX0073 / IT
C. (Carabus) granulatus CBX0043 / FR
C. (Archicarabus) nemoralis CBX0072 / FR
C. (Chrysocarabus) auronitens CBX0027 / FR
C. (Procrustes) coriaceus CBX0042 / FR
C. (Hygrocarabus) nodulosus nodulosus CBX0190 / AT
C. (Hygrocarabus) nodulosus nodulosus CBX0192 / IT
C. (Hygrocarabus) nodulosus nodulosus CBX0019 / FR
C. (Hygrocarabus) variolosus variolosus CBX0182 / RO
C. (Hygrocarabus) variolosus variolosus CBX0193 / SK
C. (Heterocarabus) marietti muchei CBX0101 / TK
C. (Chaetocarabus) arcadicus arcadicus CBX0211 / GR
C. (Chaetocarabus) arcadicus arcadicus CBX0079 / GR
C. (Chaetocarabus) intricatus lefebvrei CBX0178 / IT
C. (Chaetocarabus) intricatus intricatus CBX0078 / IT
C. (Chaetocarabus) intricatus intricatus CBX0195 / FR
C. (Platycarabus) cychroides CBX0203 / IT
C. (Platycarabus) cychroides CBX0170 / IT
C. (Platycarabus) cychroides CBX0080 / IT
C. (Platycarabus) cychroides CBX0081 / IT
C. (Platycarabus) depressus lucens CBX0207 / IT
C. (Platycarabus) depressus lucens CBX0110 / IT
C. (Platycarabus) depressus lucens CBX0183 / IT
C. (Platycarabus) depressus depressus CBX0205 / CH
C. (Platycarabus) depressus depressus CBX0111 / CH
C. (Platycarabus) depressus depressus CBX0206 / CH
C. (Platycarabus) depressus depressus CBX0085 / IT
C. (Platycarabus) depressus depressus CBX0088 /FR
C. (Platycarabus) depressus depressus CBX0108 FR
C. (Platycarabus) creutzeri baldensis CBX0092 / IT
C. (Platycarabus) creutzeri rinaldoi CBX0119 / IT
C. (Platycarabus) creutzeri viridimicans CBX0120 / AT
C. (Platycarabus) creutzeri kircheri CBX0106 / AT
C. (Platycarabus) fabricii fabricii CBX0094 / AT
C. (Platycarabus) fabricii fabricii CBX0112 / AT
C. (Platycarabus) fabricii fabricii (PT ssp. fassati) CBX0095 / SK
C. (Platycarabus) fabricii fabricii CBX0171 / SK
C. (Platycarabus) fabricii malachiticus CBX0172 / RO
C. (Platycarabus) fabricii fabricii CBX0093 / CH
C. (Platycarabus) fabricii fabricii CBX0113 / CH
C. (Platycarabus) irregularis montandoni CBX0166 / SK
C. (Platycarabus) irregularis montandoni CBX0165 / RO
C. (Platycarabus) irregularis bucephalus CBX0098 / SI
C. (Platycarabus) irregularis bucephalus CBX0168 / SI
C. (Platycarabus) irregularis irregularis CBX0115 / AT
C. (Platycarabus) irregularis irregularis CBX0194 / HU
C. (Platycarabus) irregularis irregularis CBX0099 / FR
C. (Platycarabus) irregularis irregularis CBX0208 / CH
C. (Platycarabus) irregularis irregularis CBX0020 / FR
100/100
62/73
100/76
78.8/57
100/76
100/100
100/100
100/100
99.5/97
0.008

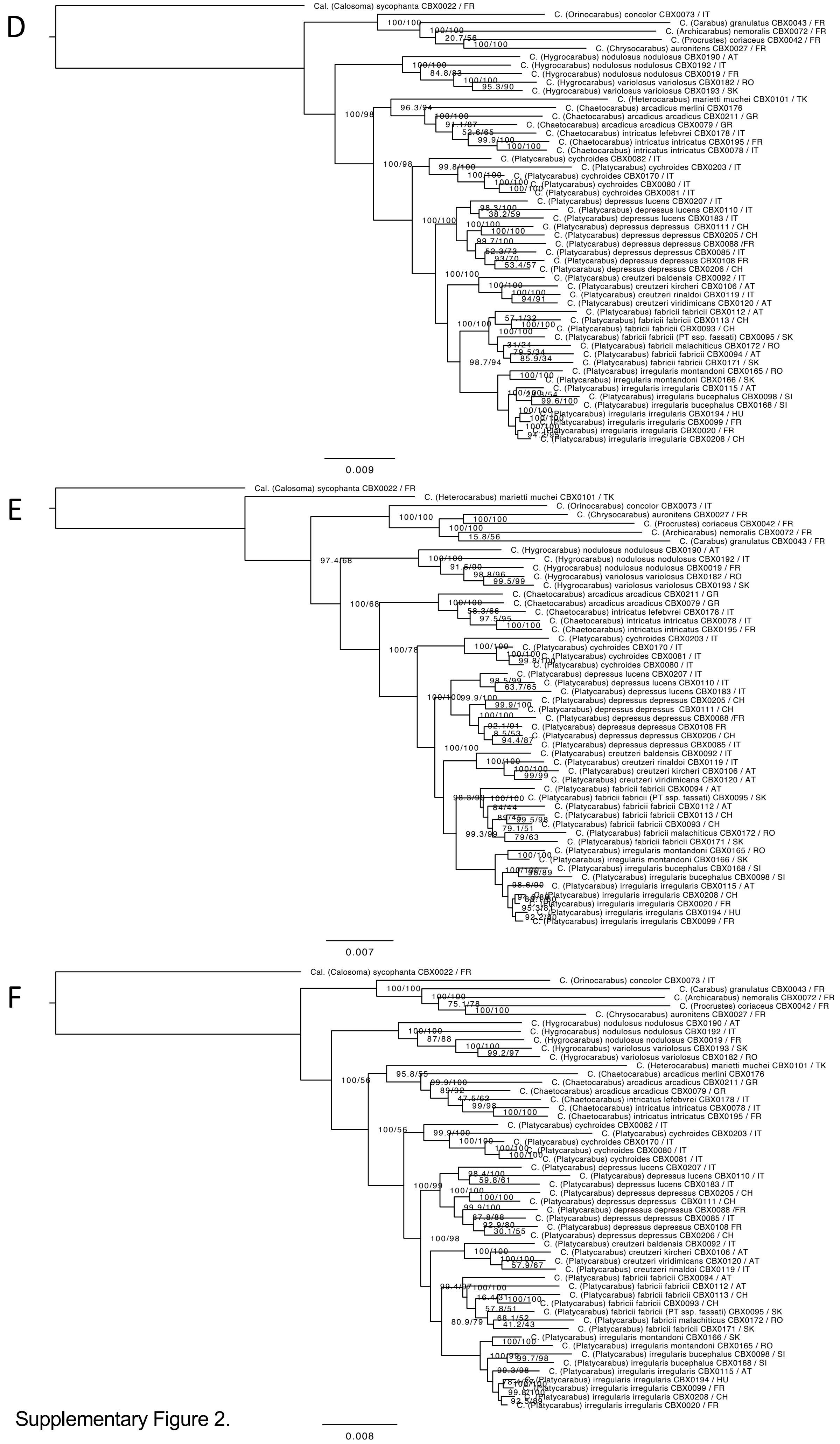


Supplementary Figure 2.

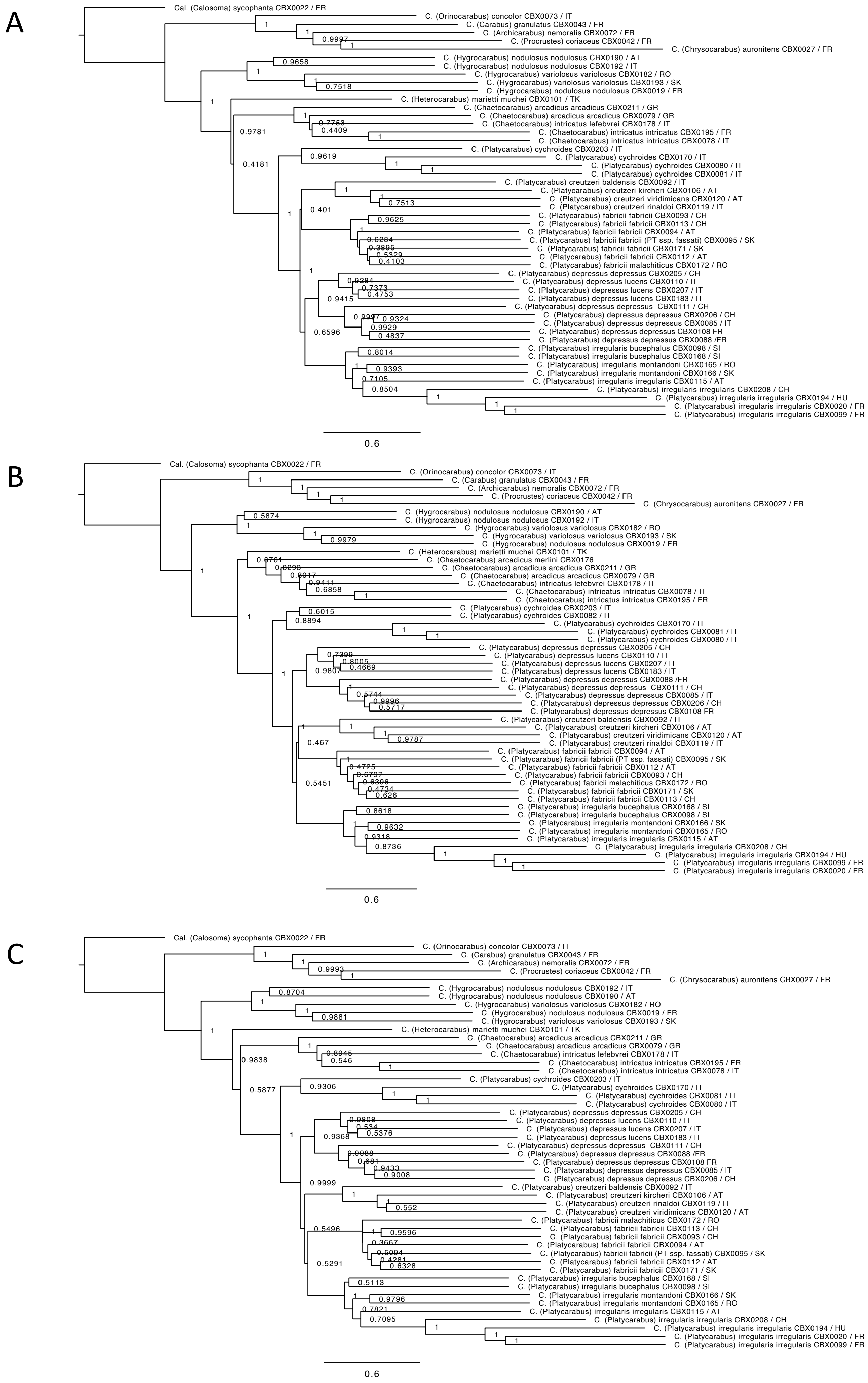
A
Cal. (Calosoma) sycophanta CBX0022 / FR
C. (Orinocarabus) concolor CBX0073 / IT
C. (Carabus) granulatus CBX0043 / FR
C. (Archicarabus) nemoralis CBX0072 / FR
C. (Procrustes) coriaceus CBX0042 / FR
C. (Chrysocarabus) auronitens CBX0027 / FR
C. (Hygrocarabus) nodulosus nodulosus CBX0190 / AT
C. (Hygrocarabus) nodulosus nodulosus CBX0192 / IT
C. (Hygrocarabus) variolosus variolosus CBX0182 / RO
C. (Hygrocarabus) variolosus variolosus CBX0193 / SK
C. (Hygrocarabus) nodulosus nodulosus CBX0019 / FR
C. (Heterocarabus) marietti muchei CBX0101 / TK
C. (Chaetocarabus) arcadicus arcadicus CBX0211 / GR
C. (Chaetocarabus) arcadicus arcadicus CBX0079 / GR
C. (Chaetocarabus) intricatus lefebvrei CBX0178 / IT
C. (Chaetocarabus) intricatus intricatus CBX0195 / FR
C. (Chaetocarabus) intricatus intricatus CBX0078 / IT
C. (Platycarabus) cychroides CBX0203 / IT
C. (Platycarabus) cychroides CBX0170 / IT
C. (Platycarabus) cychroides CBX0080 / IT
C. (Platycarabus) cychroides CBX0081 / IT
C. (Platycarabus) creutzeri baldensis CBX0092 / IT
C. (Platycarabus) creutzeri kircheri CBX0106 / AT
C. (Platycarabus) creutzeri viridimicans CBX0120 / AT
C. (Platycarabus) creutzeri rinaldoi CBX0119 / IT
C. (Platycarabus) fabricii fabricii CBX0093 / CH
C. (Platycarabus) fabricii fabricii CBX0113 / CH
C. (Platycarabus) fabricii fabricii CBX0094 / AT
C. (Platycarabus) fabricii fabricii (PT ssp. fassati) CBX0095 / SK
C. (Platycarabus) fabricii fabricii CBX0171 / SK
C. (Platycarabus) fabricii fabricii CBX0112 / AT
C. (Platycarabus) fabricii malachiticus CBX0172 / RO
C. (Platycarabus) depressus depressus CBX0205 / CH
C. (Platycarabus) depressus lucens CBX0110 / IT
C. (Platycarabus) depressus lucens CBX0207 / IT
C. (Platycarabus) depressus lucens CBX0183 / IT
C. (Platycarabus) depressus depressus CBX0111 / CH
C. (Platycarabus) depressus depressus CBX0206 / CH
C. (Platycarabus) depressus depressus CBX0085 / IT
C. (Platycarabus) depressus depressus CBX0108 FR
C. (Platycarabus) depressus depressus CBX0088 /FR
C. (Platycarabus) irregularis bucephalus CBX0098 / SI
C. (Platycarabus) irregularis bucephalus CBX0168 / SI
C. (Platycarabus) irregularis montandoni CBX0165 / RO
C. (Platycarabus) irregularis montandoni CBX0166 / SK
C. (Platycarabus) irregularis irregularis CBX0115 / AT
C. (Platycarabus) irregularis irregularis CBX0208 / CH
C. (Platycarabus) irregularis irregularis CBX0194 / HU
C. (Platycarabus) irregularis irregularis CBX0020 / FR
C. (Platycarabus) irregularis irregularis CBX0099 / FR
0.6
B
Cal. (Calosoma) sycophanta CBX0022 / FR
C. (Orinocarabus) concolor CBX0073 / IT
C. (Carabus) granulatus CBX0043 / FR
C. (Archicarabus) nemoralis CBX0072 / FR
C. (Procrustes) coriaceus CBX0042 / FR
C. (Chrysocarabus) auronitens CBX0027 / FR
C. (Hygrocarabus) nodulosus nodulosus CBX0190 / AT
C. (Hygrocarabus) nodulosus nodulosus CBX0192 / IT
C. (Hygrocarabus) variolosus variolosus CBX0182 / RO
C. (Hygrocarabus) variolosus variolosus CBX0193 / SK
C. (Hygrocarabus) nodulosus nodulosus CBX0019 / FR
C. (Heterocarabus) marietti muchei CBX0101 / TK
C. (Chaetocarabus) arcadicus merlini CBX0176
C. (Chaetocarabus) arcadicus arcadicus CBX0211 / GR
C. (Chaetocarabus) arcadicus arcadicus CBX0079 / GR
C. (Chaetocarabus) intricatus lefebvrei CBX0178 / IT
C. (Chaetocarabus) intricatus intricatus CBX0078 / IT
C. (Chaetocarabus) intricatus intricatus CBX0195 / FR
C. (Platycarabus) cychroides CBX0203 / IT
C. (Platycarabus) cychroides CBX0082 / IT
C. (Platycarabus) cychroides CBX0170 / IT
C. (Platycarabus) cychroides CBX0081 / IT
C. (Platycarabus) cychroides CBX0080 / IT
C. (Platycarabus) depressus depressus CBX0205 / CH
C. (Platycarabus) depressus lucens CBX0110 / IT
C. (Platycarabus) depressus lucens CBX0207 / IT
C. (Platycarabus) depressus lucens CBX0183 / IT
C. (Platycarabus) depressus depressus CBX0088 /FR
C. (Platycarabus) depressus depressus CBX0111 / CH
C. (Platycarabus) depressus depressus CBX0085 / IT
C. (Platycarabus) depressus depressus CBX0206 / CH
C. (Platycarabus) depressus depressus CBX0108 FR
C. (Platycarabus) creutzeri baldensis CBX0092 / IT
C. (Platycarabus) creutzeri kircheri CBX0106 / AT
C. (Platycarabus) creutzeri viridimicans CBX0120 / AT
C. (Platycarabus) creutzeri rinaldoi CBX0119 / IT
C. (Platycarabus) fabricii fabricii CBX0094 / AT
C. (Platycarabus) fabricii fabricii (PT ssp. fassati) CBX0095 / SK
C. (Platycarabus) fabricii fabricii CBX0112 / AT
C. (Platycarabus) fabricii fabricii CBX0093 / CH
C. (Platycarabus) fabricii malachiticus CBX0172 / RO
C. (Platycarabus) fabricii fabricii CBX0171 / SK
C. (Platycarabus) fabricii fabricii CBX0113 / CH
C. (Platycarabus) irregularis bucephalus CBX0168 / SI
C. (Platycarabus) irregularis bucephalus CBX0098 / SI
C. (Platycarabus) irregularis montandoni CBX0166 / SK
C. (Platycarabus) irregularis montandoni CBX0165 / RO
C. (Platycarabus) irregularis irregularis CBX0115 / AT
C. (Platycarabus) irregularis irregularis CBX0208 / CH
C. (Platycarabus) irregularis irregularis CBX0194 / HU
C. (Platycarabus) irregularis irregularis CBX0099 / FR
C. (Platycarabus) irregularis irregularis CBX0020 / FR
0.6
C
Cal. (Calosoma) sycophanta CBX0022 / FR
C. (Orinocarabus) concolor CBX0073 / IT
C. (Carabus) granulatus CBX0043 / FR
C. (Archicarabus) nemoralis CBX0072 / FR
C. (Procrustes) coriaceus CBX0042 / FR
C. (Chrysocarabus) auronitens CBX0027 / FR
C. (Hygrocarabus) nodulosus nodulosus CBX0192 / IT
C. (Hygrocarabus) nodulosus nodulosus CBX0190 / AT
C. (Hygrocarabus) variolosus variolosus CBX0182 / RO
C. (Hygrocarabus) nodulosus nodulosus CBX0019 / FR
C. (Hygrocarabus) variolosus variolosus CBX0193 / SK
C. (Heterocarabus) marietti muchei CBX0101 / TK
C. (Chaetocarabus) arcadicus arcadicus CBX0211 / GR
C. (Chaetocarabus) arcadicus arcadicus CBX0079 / GR
C. (Chaetocarabus) intricatus lefebvrei CBX0178 / IT
C. (Chaetocarabus) intricatus intricatus CBX0195 / FR
C. (Chaetocarabus) intricatus intricatus CBX0078 / IT
C. (Platycarabus) cychroides CBX0203 / IT
C. (Platycarabus) cychroides CBX0170 / IT
C. (Platycarabus) cychroides CBX0081 / IT
C. (Platycarabus) cychroides CBX0080 / IT
C. (Platycarabus) depressus depressus CBX0205 / CH
C. (Platycarabus) depressus lucens CBX0110 / IT
C. (Platycarabus) depressus lucens CBX0207 / IT
C. (Platycarabus) depressus lucens CBX0183 / IT
C. (Platycarabus) depressus depressus CBX0111 / CH
C. (Platycarabus) depressus depressus CBX0088 /FR
C. (Platycarabus) depressus depressus CBX0108 FR
C. (Platycarabus) depressus depressus CBX0085 / IT
C. (Platycarabus) depressus depressus CBX0206 / CH
C. (Platycarabus) creutzeri baldensis CBX0092 / IT
C. (Platycarabus) creutzeri kircheri CBX0106 / AT
C. (Platycarabus) creutzeri rinaldoi CBX0119 / IT
C. (Platycarabus) creutzeri viridimicans CBX0120 / AT
C. (Platycarabus) fabricii malachiticus CBX0172 / RO
C. (Platycarabus) fabricii fabricii CBX0113 / CH
C. (Platycarabus) fabricii fabricii CBX0093 / CH
C. (Platycarabus) fabricii fabricii CBX0094 / AT
C. (Platycarabus) fabricii fabricii (PT ssp. fassati) CBX0095 / SK
C. (Platycarabus) fabricii fabricii CBX0112 / AT
C. (Platycarabus) fabricii fabricii CBX0171 / SK
C. (Platycarabus) irregularis bucephalus CBX0168 / SI
C. (Platycarabus) irregularis bucephalus CBX0098 / SI
C. (Platycarabus) irregularis montandoni CBX0166 / SK
C. (Platycarabus) irregularis montandoni CBX0165 / RO
C. (Platycarabus) irregularis irregularis CBX0115 / AT
C. (Platycarabus) irregularis irregularis CBX0208 / CH
C. (Platycarabus) irregularis irregularis CBX0194 / HU
C. (Platycarabus) irregularis irregularis CBX0020 / FR
C. (Platycarabus) irregularis irregularis CBX0099 / FR
0.6

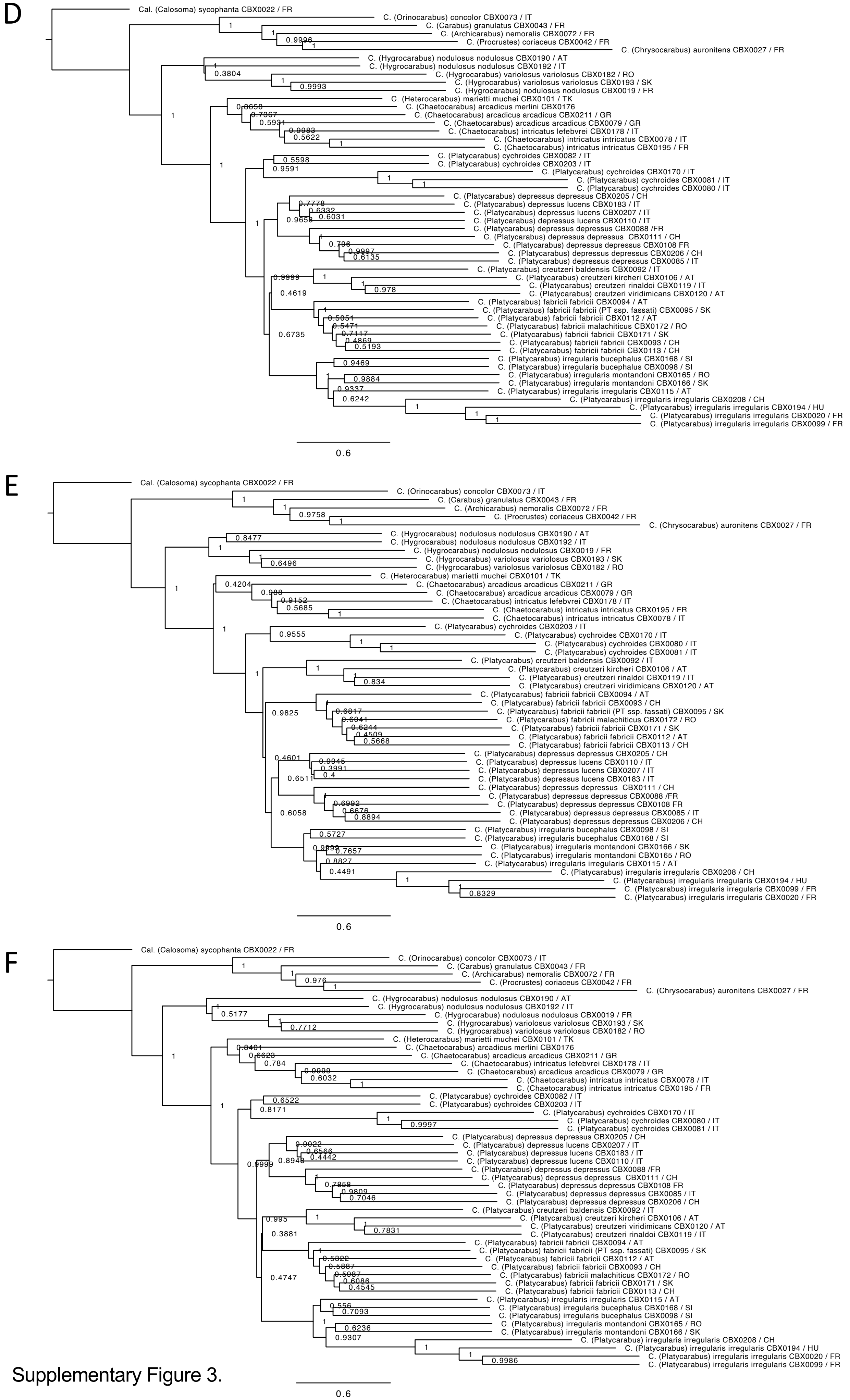


Supplementary Figure 3.

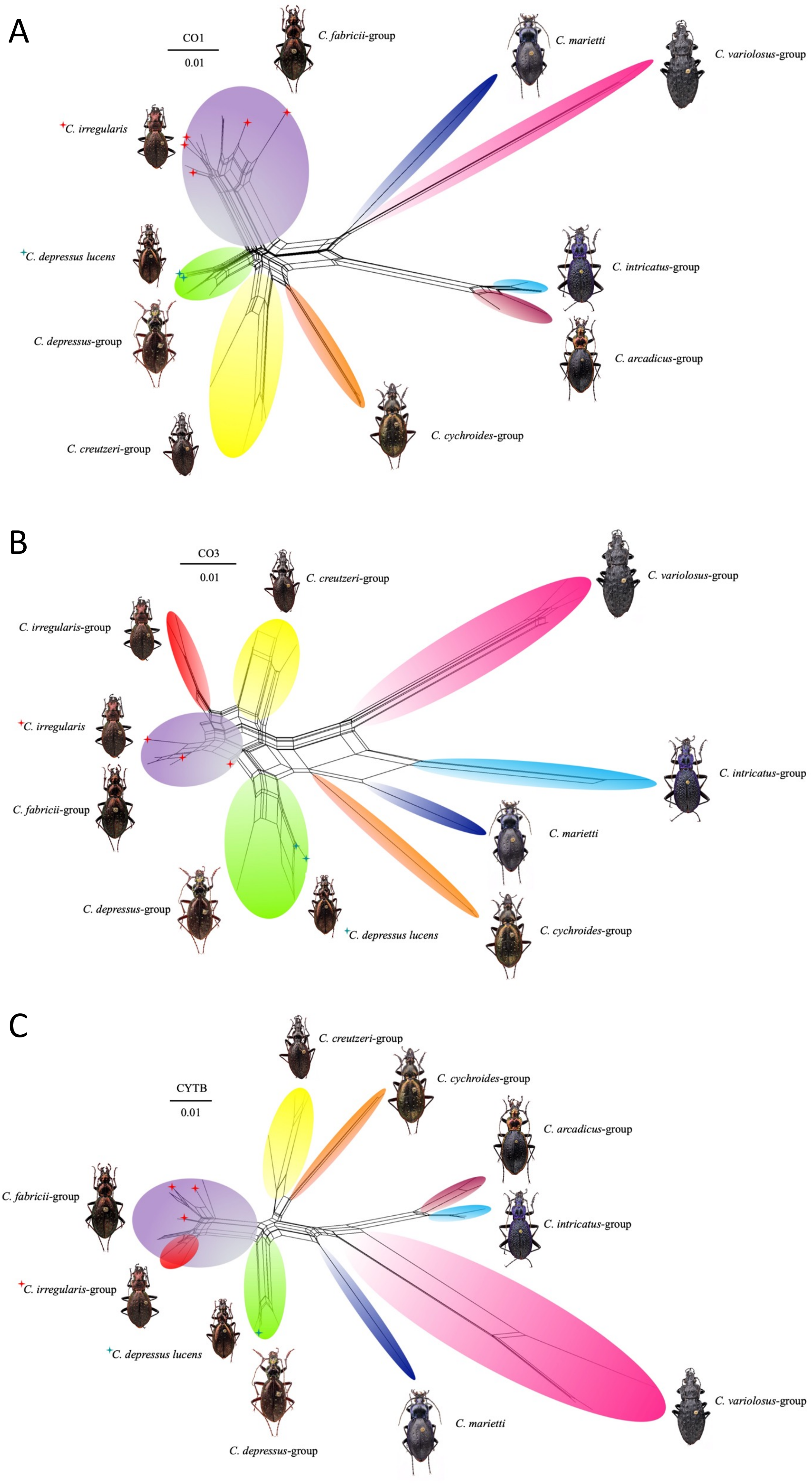


Supplementary Figure 4.

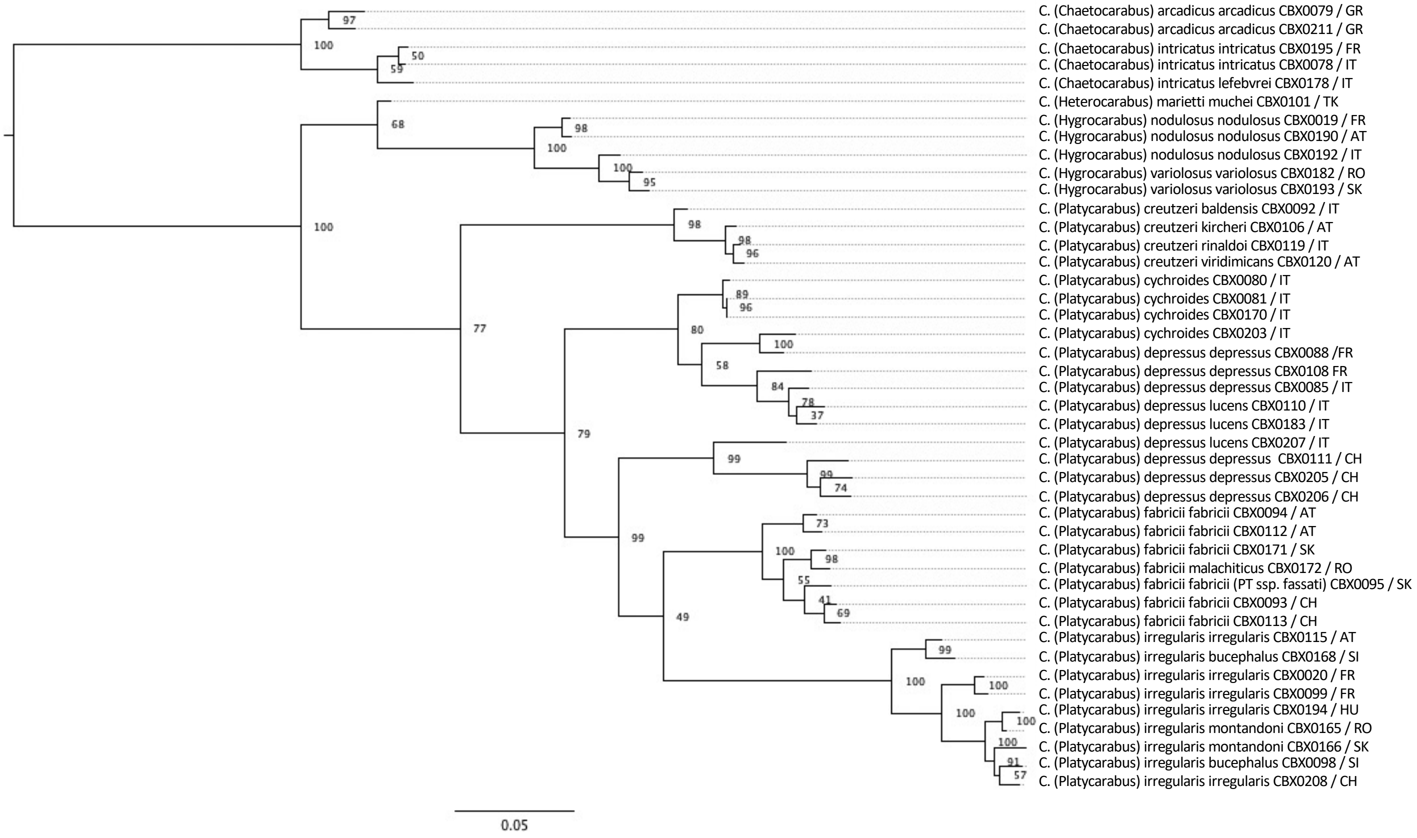


Supplementary Figure 5.

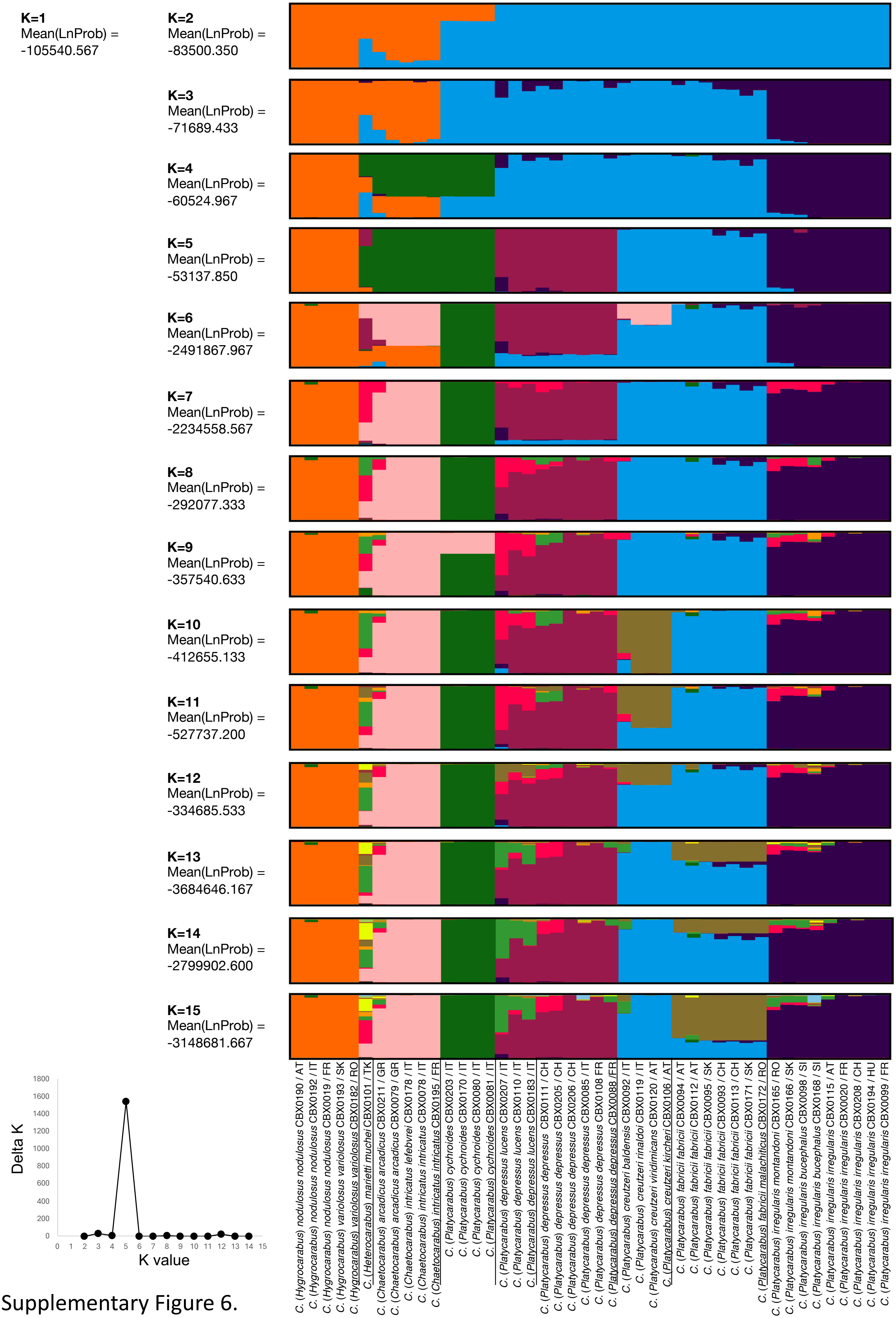


Supplementary Figure 6.

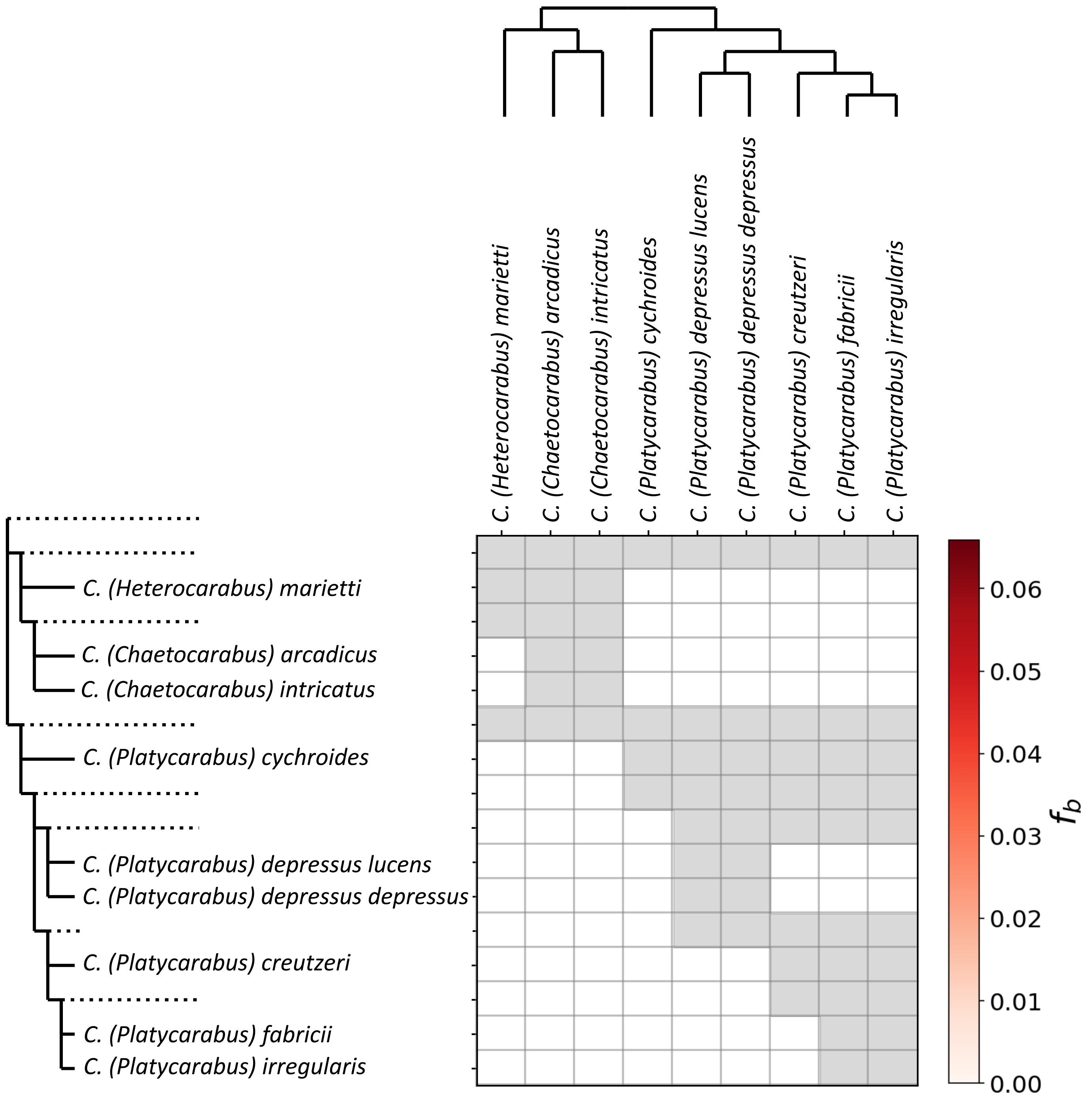


Supplementary Figure 7.
